# Metformin boosts mitochondria and neurogenesis via AMPK/mTOR/SIRT3 in POLG mutant organoids

**DOI:** 10.1101/2025.06.08.658480

**Authors:** Anbin Chen, Sebastian Edgar Schmidke, Cecilie Katrin Kristiansen, Kristina Xiao Liang

**Affiliations:** Department of Neurosurgery, Xinhua Hospital Affiliated to Shanghai Jiaotong University School of Medicine, Shanghai, 20092, China; Department of Biomedicine (IBM), Faculty of Medicine, University of Bergen, Bergen, 5009, Norway; Bergen Center of Medicine Stem Cell Research (BCMS), Faculty of Medicine, University of Bergen, Bergen, 5009, Norway; Department of Clinical Medicine (K1), Faculty of Medicine, University of Bergen, Bergen, 5009, Norway

**Author notes:** **Corresponding author** Kristina Xiao Liang Department of Biomedicine, University of Bergen, Bergen, Norway, PO Box 5009, Bergen, Norway.

**Keywords:** POLG, Cortical organoid, AMPK signaling, Mitochondrial biogenesis, SIRT3, Neurogenesis, Cortical organoids

## Abstract

**Background:** Mutations in the *POLG* gene, encoding the catalytic subunit of mitochondrial DNA polymerase gamma, are the most common cause of mitochondrial diseases affecting the central nervous system. These mutations frequently result in neurodevelopmental disorders, yet the cellular and molecular mechanisms underlying POLG related encephalopathies remain poorly understood. In particular, how POLG mutations affect mitochondrial function, neural progenitor behavior, and early neurogenesis in the developing human brain has not been fully elucidated.

**Methods:** To investigate the impact of *POLG* mutations on human neurodevelopment, we generated 3D cortical brain organoids from induced pluripotent stem cells (iPSCs) derived from a patient carrying compound heterozygous *POLG* mutations (A467T/W748S). Organoid development was monitored using immunohistochemistry, transmission electron microscopy, and live-cell mitochondrial assays. Single-cell RNA sequencing (scRNA-seq) was performed to profile cellular diversity and transcriptional changes. Organoids were treated with metformin, a known mitochondrial modulator, and mitochondrial function was assessed by measuring membrane potential (TMRE), ATP production, and mtDNA copy number. Western blotting and immunofluorescence were used to investigate AMPK–SIRT3–mTOR signaling and markers of mitochondrial dynamics and mitophagy. Statistical analyses were performed using unpaired t-tests or ANOVA, with p-values < 0.05 considered significant.

**Results:** *POLG* mutant cortical organoids exhibited impaired neural differentiation, with expansion of stress-associated neural progenitors and reduced neuronal populations. scRNA seq analysis revealed transcriptional signatures of oxidative stress, mitochondrial dysfunction, and suppressed neurogenesis, particularly in a specific neural progenitor subpopulation. Metformin treatment significantly improved mitochondrial membrane potential, ATP output, and mtDNA copy number in POLG organoids. It also promoted neuronal differentiation and reduced reactive progenitor states. Mechanistically, metformin activated AMPK and SIRT3, inhibited mTOR, enhanced expression of mitochondrial fusion and biogenesis markers (OPA1, PGC-1α), and increased autophagic and mitophagic activity (LC3B, BNIP3).

**Conclusions:** Our study demonstrates that *POLG* mutations disrupt early cortical development by impairing mitochondrial function and skewing progenitor fate. Metformin mitigates these effects by restoring mitochondrial homeostasis and promoting neurogenesis through the AMPK/SIRT3/mTOR axis. These findings offer mechanistic insights into POLG-related encephalopathy and support metformin as a candidate for therapeutic intervention in mitochondrial neurodevelopmental disorders.

## Introduction

Mitochondrial dysfunction is a well-documented hallmark of neurodegenerative diseases, with a particular emphasis on conditions involving mutations in mitochondrial DNA (mtDNA) polymerase gamma (POLG). *POLG* mutations lead to impaired mtDNA replication and repair, compromising mitochondrial integrity and function, which are essential for energy production, particularly in the brain and other high-demand tissues [1, 2]. Patients with *POLG* mutations exhibit progressive neurological symptoms, such as encephalopathy, myopathy, and seizures [1], reflecting the critical role of mitochondrial health in maintaining neural function. These mutations destabilize mitochondrial homeostasis, reduce adenosine triphosphate (ATP) synthesis, elevate oxidative stress, collectively accelerating neurodegenerative processes [3, 4].

Therapeutic strategies targeting mitochondrial dysfunction are urgently needed, and recent attention has turned to pharmacological agents that can modulate mitochondrial biogenesis and function. Metformin, a widely used antidiabetic medication, has garnered interest in mitochondrial disease research due to its activation of AMP-activated protein kinase (AMPK), a master regulator of metabolism and mitochondrial homeostasis homeostasis [5, 6]. Activation of AMPK by metformin has been shown to improve mitochondrial respiration and enhance mitochondrial function [7]. Additionally, metformin’s activation of AMPK contributes to its various therapeutic effects, including metabolic regulation and potential neuroprotective benefits [8, 9].

Recent advances in patient-specific, induced pluripotent stem cell (iPSC)-derived models have significantly advanced our understanding of mitochondrial diseases [4, 6, 10, 11]. These models allow the study of mitochondrial dynamics and neurogenesis within a personalized genetic context, providing deeper insights into the cell-specific impacts of *POLG* mutations. For instance, iPSC-derived cortical organoids [11] and neural progenitors (NPCs) [4] from patients carrying *POLG* mutations offer a closer approximation to human neural development and function, making them highly suitable for investigating the effects of potential therapeutic agents like metformin. Leveraging these advanced models could elucidate metformin’s impact on mitochondrial function and neural cell health in *POLG*-mutant backgrounds.

In this study, we sought to assess whether metformin could restore mitochondrial respiratory function and alleviate neural deficits in POLG-mutant cells derived from CP2A patient carrying compound heterozygous c.1399 G > A/c.2243 G > C, p.A467T/W748S). Using a comprehensive suite of patient-derived neural models—including cortical organoids, NPCs, mature neurons, and neural spheres, we aimed to investigate metformin’s potential to enhance mitochondrial function, reduce oxidative stress, and promote cellular viability within a patient-specific setting. We hypothesized that metformin would improve mitochondrial health in *POLG*-mutant cells, potentially opening a therapeutic avenue for mitochondrial dysfunction-associated neurodegenerative diseases.

## Materials and Methods

### iPSC and NPC generation

iPSCs were generated using virus-vector reprogramming method from skin fibroblasts of healthy donors, and cells were cultured in Essential 8 medium and maintained as previously described [4]. NPCs were derived from the iPSCs using a dual SMAD inhibition protocol. iPSCs were dissociated with gentle cell dissociation reagent and plated on Matrigel-coated plates in neural induction medium containing N2 supplement (Thermo Fisher Scientific, 17502048), B27 supplement minus vitamin A (Thermo Fisher Scientific, 12587012), 10 μM SB431542 (Tocris Bioscience, 1614; TGF-β inhibitor), and 1 μM Dorsomorphin (Tocris Bioscience, 3093; BMP inhibitor). The medium was changed every other day. After 10 days of induction, rosette structures typical of neural progenitors were manually selected and expanded in StemPro NPC medium supplemented with EGF and FGF2 (Thermo Fisher Scientific, A1050901).

### Cortical organoid generation

Cortical organoids were generated following previously established protocols [11]. Feeder-free iPSCs were cultured in E8 medium for 7 days before differentiation. Once iPSCs reached 70% confluency, they were dissociated with Accutase (Sigma-Aldrich, A6964) for 10 minutes at 37°C, followed by centrifugation. Approximately 9,000 cells were seeded into 96-well ultra-low attachment plates in neural induction medium with 50 μM ROCK inhibitor (Millipore, SCM075). Differentiation was initiated using dual SMAD and WNT inhibitors. Embryoid bodies formed within 24 hours, and the medium was refreshed on Days 2, 4, 6, and 8. On Day 10, organoids were transferred to 6-well ultra-low attachment plates with neural differentiation medium without vitamin A and placed on an orbital shaker. From Day 18, vitamin A (Invitrogen, 17504044) and BDNF (R&D System, 248-BD) were added to support maturation. The medium was replaced every 3–4 days to sustain growth and maturation.

### Metformin treatment

NPCs were exposed to metformin (Sigma-Aldrich, 317240) for a duration of 5 days as part of the phenotype rescue experiments. For cortical organoids, the same concentration of metformin was introduced on the sixth day of differentiation, with treatment continuing over a two-month period. The culture medium was renewed every 48 hours to maintain optimal conditions. Control groups, including untreated NPCs and organoids, were cultured under the same conditions, with dimethyl sulfoxide (DMSO, Sigma-Aldrich, 153087) added to the medium in equivalent volumes to serve as the vehicle control.

### Immunofluorescence Staining and Imaging

For NPC staining, cells cultured on glass coverslips were fixed with 4% paraformaldehyde (PFA; Thermo Fisher Scientific, 28908) in PBS for 15 minutes at room temperature, followed by two PBS washes. Permeabilization and blocking were performed using 10% goat serum (Sigma-Aldrich, 50197Z) and 0.3% Triton X-100 (Sigma-Aldrich, 9036-19-5) in PBS for 1 hour at room temperature. Primary antibodies diluted in blocking buffer were applied overnight at 4°C. After PBS washes, secondary antibodies and Hoechst 33324 nuclear stain were added and incubated for 1 hour at room temperature in the dark. Coverslips were mounted with Fluoromount-G mounting medium (SouthernBiotech, 0100-20), sealed with 1.5 mm coverslips, and cured at room temperature in the dark for 24 hours before imaging with a Leica TCS SP8 STED 3X confocal microscope (Leica Microsystems).

For cortical organoid staining, intact organoids were transferred onto Superfrost™ adhesion slides (Thermo Fisher Scientific, J1800AMNZ) using a cut-tip pipette. After air-drying, organoids were fixed with 4% PFA for 30 minutes at room temperature and washed twice with PBS. Blocking and permeabilization were carried out using 10% goat serum and 0.1% Triton X-100 in PBS for 2 hours at room temperature. Primary antibodies diluted in blocking buffer were incubated overnight at 4°C. After PBS washes, secondary antibodies and Hoechst stain were applied and incubated for 48 hours at 4°C in the dark. Organoids were mounted using Fluoromount-G, covered with 1.5 mm coverslips, cured at room temperature for 24 hours, and stored at –20°C until imaging. Imaging was performed using the Leica TCS SP8 STED 3X confocal microscope.

Primary antibodies used included: TFAM (mouse, 1:1000; Abcam, ab119684); β-tubulin III (TUJ1) (mouse, 1:1000; Abcam, ab78078); NDUFB10 (rabbit, 1:350; Abcam, ab196019); TOMM20 (mouse, 1:350; Abcam, ab56783); GFAP (chicken, 1:500; Abcam, ab4674); NeuN (rabbit, 1:500; Cell Signaling Technology, 24307S); CTIP2 (rat, 1:500; Abcam, ab18465); SOX2 (rabbit, 1:100; Abcam, ab97959). Secondary antibodies conjugated to Alexa Fluor dyes (Invitrogen, A11008, A21449, A11012, A21141, A11042, A21236) were used at 1:800 dilution. Coverslips were mounted using ProLong Diamond Antifade Mountant (Thermo Fisher Scientific, P36965).

Immunofluorescence images were quantitatively analyzed using ImageJ software (Version 1.52a; NIH, USA). Six to ten regions within cortical layers were randomly selected for measurement. Single-channel images were converted to 8-bit grayscale, and default threshold settings were used to minimize selection bias. Fluorescence intensity was calculated as the mean gray value (Mean = Integrated Density/Area). Quantitative data were analyzed and plotted using GraphPad Prism 8.0.2 software (GraphPad Software, Inc.).

### ScRNA-seq and data analysis

#### Organoid dissociation and single-cell isolation

Brain organoids were harvested, rinsed with 1x PBS, and cut into 1–2 mm fragments using ophthalmic scissors. The fragments were digested in Cell Live Tissue Dissociation Solution at 37°C for 15 minutes with agitation. The resulting cell suspension was filtered through a 40-μm strainer, centrifuged at 350 × g for 5 minutes at 4°C, and resuspended in PBS. Cell viability and concentration were assessed using Trypan Blue staining and a hemocytometer.

#### scRNA-Seq library preparation and sequencing

Single-cell RNA sequencing (scRNA-seq) libraries were prepared using the GEXSCOPE Library Kit, following the manufacturer’s protocol. The cell suspension was loaded onto a microfluidic chip, capturing 6000 cells. After mRNA capture and cDNA amplification, sequencing libraries were constructed and analyzed using an Agilent Fragment Analyzer. The final libraries were sequenced on an Illumina NovaSeq 6000 platform with paired-end 2 × 150 bp, yielding 90 GB per library.

#### Tranpcriptome data processing and quality control

Raw sequencing data was processed using CeleScope v1.3.0, with reads aligned to the human reference genome (STAR aligner). Gene annotations were derived from Ensembl 92, and gene counts were generated using featureCounts. Quality control was performed using Scanpy, filtering out cells with >20% mitochondrial reads, potential doublets (>5000 genes), and low-quality cells (<200 genes detected).

#### Sample integration, dimensionality reduction, and clustering

Datasets were combined using and data’s concatenate function, normalized to 10,000 counts per cell, and analyzed for highly variable genes. Principal component analysis (PCA) was performed, selecting the top 17 principal components. The UMAP algorithm was applied for dimensionality reduction, and cell clusters were identified using the Leiden algorithm (resolution = 0.5).

#### Cell type annotation and differential gene expression analysis

Cell clusters were annotated using scMRMA, with marker genes identified via PanglaoDB and validated using Fisher’s enrichment analysis. Differentially expressed genes (DEGs) were identified using Wilcoxon rank-sum test (Bonferroni correction), with a p-value < 0.05 considered significant.

#### Gene enrichment and pathway analysis

Gene enrichment was performed using gseapy, querying databases such as GO_Biological_Process, Reactome, KEGG, and KEA.

#### Statistical analysis

The data were presented as mean ± standard deviation for a sample size of at least three. Normality of the data was assessed using the Shapiro-Wilk test, and potential outliers were identified via the ROUT method. For non-normally distributed variables, the Mann-Whitney U-test was employed to determine statistical significance, whereas a two-sided Student’s T-test was used for normally distributed variables. All statistical analyses were conducted using GraphPad Prism 8.0.2, with significance defined as p ≤ 0.05.

## Results

### Establishment of 3D cortical organoids model from iPSCs

We generated iPSCs from skin fibroblasts of a patient with *POLG* mutations and two neurologically healthy individual controls, following our previously established protocols [4, 10]. The patient carried compound heterozygous mutations c.1399 G > A (p.A467T) and c.2243 G > C (p.W748S) (CP2A) and exhibited symptoms characteristic of POLG-related disorders, including progressive spinocerebellar ataxia, extraocular myopathy, migraine-like headaches, and peripheral neuropath [2].

We further differentiated POLG patient iPSCs and control iPSCs into 3D cortical organoids, following a stepwise differentiation protocol [11], through three key stages: neural induction, differentiation, and maturation (Fig. 1A). The differentiation process was monitored over time, revealing progressive morphological development and structural complexity. On Day 1-2, iPSCs formed compact and tightly clustered spheroids, which further increased in size and complexity as differentiation progressed (Day 18). By Day 30, the organoids exhibited defined cortical-like structures with distinct layers (Fig. 1B).

**Figure 1.**
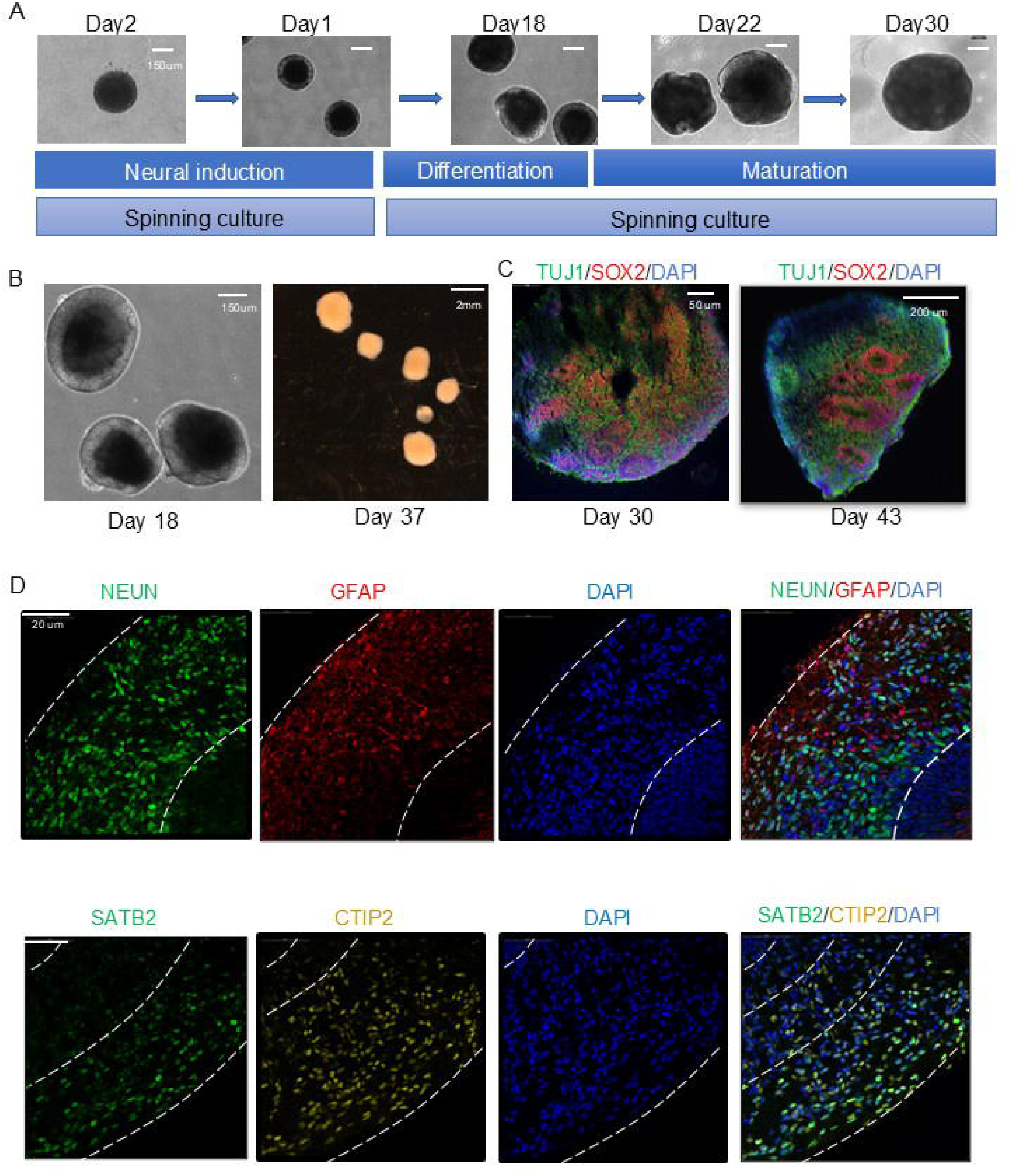
Stepwise generation and characterization of iPSC-derived 3D cortical organoids. **(A)** Schematic representation of the cortical organoid differentiation protocol. Human iPSCs were aggregated and cultured in spinning suspension from day 0, progressing through neural induction (day 1–2), differentiation (day 18), and early maturation stages (day 22–30). Representative Bright-field images show progressive morphological development of organoids over time. Scale bars: 150 μm. **(B)** Representative images of organoid morphology during early development. a, Bright-field images showing compact and spherical organoids at early stages. b, Macroscopic view of multiple organoids after differentiation. Scale bars: 2 mm. **(C)** Immunofluorescence staining of cortical organoids at day 30 showing expression of TUJ1 (green), SOX2 (red), and DAPI (blue), demonstrating co-localization of neural progenitor and early neuronal markers. Scale bars: 50 μm (left), 200 μm (right). **(D)** Immunostaining for mature cortical markers at later stages. Left panel: NEUN (green) and GFAP (red) indicate the presence of mature neurons and astrocytes, respectively, with DAPI (blue) counterstaining. Right panel: Layer-specific cortical markers SATB2 (upper layer, green) and CTIP2 (deep layer, yellow) reveal the formation of structured cortical plate-like organization. Dashed lines indicate cortical layer boundaries. Scale bars: 20 μm.

Immunostaining confirmed the presence of key neuronal markers at different stages of organoid maturation. At Day 30 and Day 43, we observed the expression of early neuronal marker TUJ1 and neural progenitor marker SOX2, indicating active neurogenesis (Fig. 1C). By Day 70, immunofluorescence staining revealed mature neuronal marker NEUN, astrocytic marker GFAP, and cortical layer-specific markers SATB2 (upper-layer neurons) and CTIP2 (deep-layer neurons), demonstrating the successful establishment of layered cortical structures in the organoids (Fig. 1D). These findings confirm that our iPSC-derived cortical organoids recapitulate key aspects of early cortical development and neuronal differentiation.

### Single-cell profiling uncovers distinct NPC subclusters and stress signatures in control and POLG cortical organoids

Previously, we reported that metformin enhances mitochondrial function in POLG astrocytes [6]. To further investigate the effects of metformin on NPCs, we treated POLG patient-derived cortical organoids with 500 µM metformin for two months and performed scRNA-seq. To examine the molecular characteristics and cellular heterogeneity of NPCs across different conditions-control organoids (Fig. 2A), POLG patient organoids (Fig. 2B), and metformin-treated patient organoids (Fig. 2C), we conducted scRNA-seq and uniform manifold approximation and projection (UMAP) analysis. The UMAP visualization illustrates the distribution of NPC subclusters (Fig. 2D) among the different sample groups: Control (green), POLG CP2A patient (blue), and POLG CP2A patient with metformin treatment (orange), allowing us to assess the impact of metformin treatment on the cellular landscape of POLG NPCs. The results demonstrate that NPCs from both patient-derived and control organoids exhibited two distinct but overlapping clustering patterns NPC g cluster and NPC f cluster, suggesting conserved developmental trajectories despite POLG mutations. However, noticeable differences in cellular distribution were observed, with patient-derived NPCs showing a broader spread across the UMAP space, potentially reflecting altered transcriptional profiles and developmental divergence. The proportion of NPCs varied across different organoid samples, as assessed by their relative abundance within the whole cell population (Fig. 2E) and the neural population (Fig. 2F). Specifically, POLG patient-derived organoids exhibited a higher proportion of NPCs compared to controls, suggesting a potential delay in differentiation. Next, we explored the significant transcriptional alterations in NPC population from POLG brain organoids compared to controls. A total of 5 genes were upregulated and 52 genes were downregulated (Fig. 2G), while a separate analysis identified 2 upregulated and 20 downregulated genes (Fig. 2H), indicating broad suppression of gene expression in POLG NPCs. Gene Ontology (GO) and Reactome pathway enrichment analyses showed significant downregulation of key molecular functions, including oxidoreductase activity, NAD-binding proteins, protein kinase regulatory subunit binding, RNA binding, and vascular endothelial growth factor receptor binding (Fig. 2I), suggesting impaired metabolic activity and neurodevelopmental regulatory mechanisms. Additionally, Reactome pathway analysis highlighted downregulation in pathways related to mitochondrial translation, oxidative stress response, chaperonin-mediated protein folding, and organelle biogenesis and maintenance (Fig. 2J), reinforcing the role of mitochondrial dysfunction in POLG NPC impairment. Notably, inflammatory pathways such as inflammasome activation and NF-κB signaling were also suppressed, suggesting an altered immune response, while dysregulation of lipid metabolism and ER stress-related pathways further supports the hypothesis of metabolic dysfunction. Collectively, these findings highlight widespread mitochondrial and metabolic impairments in POLG NPCs, potentially contributing to disrupted neuronal differentiation and maintenance in POLG-related disorders.

**Figure 2.**
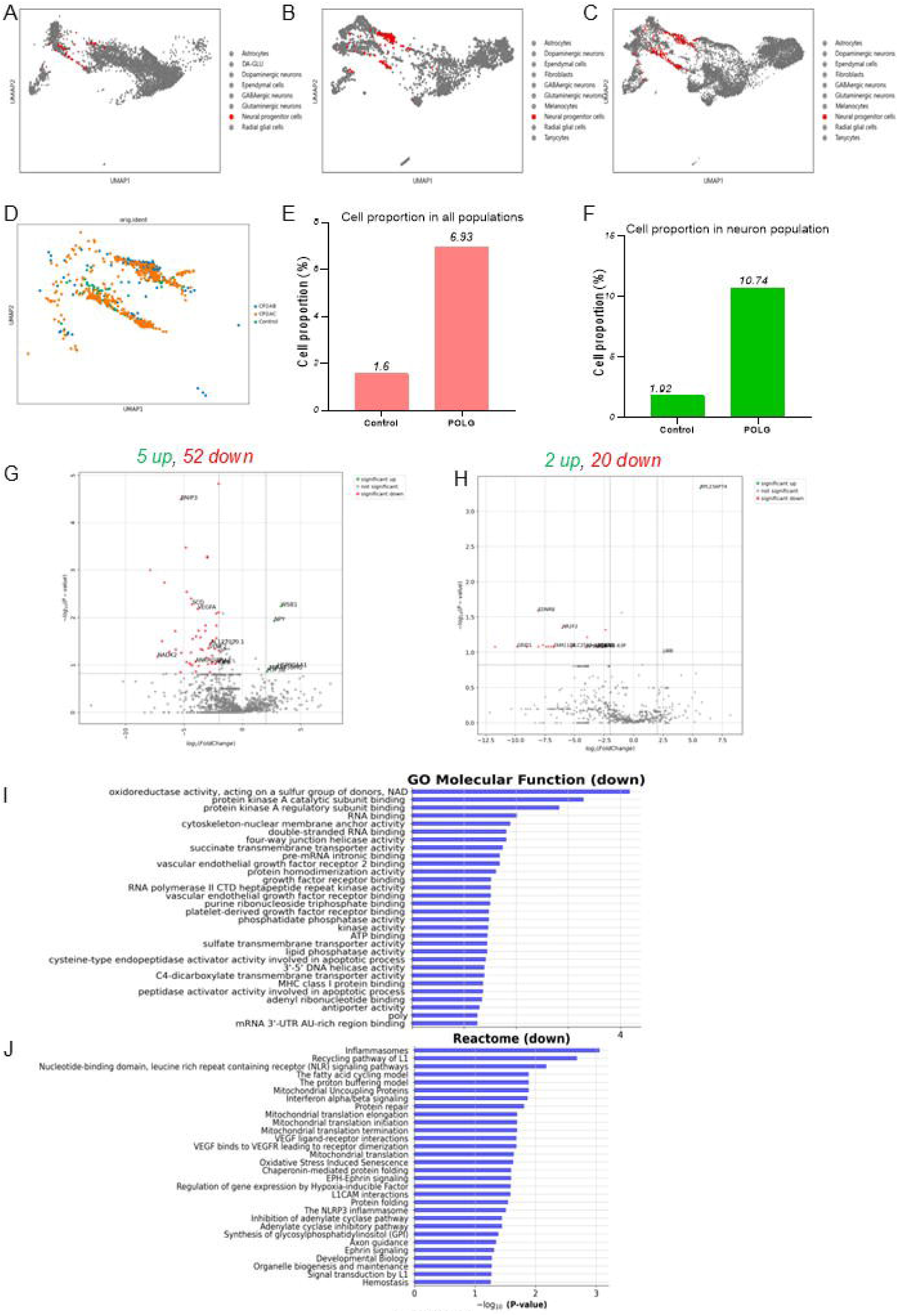
Single-cell transcriptomic profiling reveals neural progenitor cell alterations and transcriptional dysregulation in POLG patient-derived cortical organoids. (**A–C**) UMAP plots showing the spatial distribution of neural progenitor cluster g (highlighted in red) across three conditions: (A) Control organoids, (B) *POLG* mutant (CP2A) organoids, and (C) Metformin-treated POLG organoids. (I) (**D**) UMAP plot displaying integration of all three samples. Individual cells are color-coded by condition: Control (green), POLG CP2A (blue), and POLG CP2A + Metformin (orange), highlighting shifts in NPC distribution upon metformin treatment. (**E–F**) Bar graphs showing the proportion of cluster g cells relative to total cells (E) and to the neural lineage population (F) across conditions. POLG CP2A organoids exhibit increased cluster g representation, which is reduced after metformin treatment. (**G–H**) Volcano plots illustrating DEGs in NPC populations: (G) CP2A vs. Control and (H) CP2A + Metformin vs. Control. Significant upregulated genes (green), downregulated genes (red), and selected key transcripts are highlighted. POLG NPCs show broad transcriptional repression (52 down, 5 up), partially reversed by metformin (20 down, 2 up). **(I–J**) GO Molecular Function (I) and Reactome Pathway (J) enrichment analysis of downregulated genes in POLG NPCs. Functional categories related to mitochondrial translation, oxidoreductase activity, chaperonin-mediated protein folding, immune response (e.g., inflammasome, NF-κB signaling), and cytoskeletal organization are significantly suppressed, suggesting widespread metabolic and developmental impairment in POLG neural progenitors.

We further analyzed the subclusters within brain organoids derived from control individuals and POLG patients to explore how POLG mutations influence cellular heterogeneity and gene expression profiles at a finer resolution. The UMAP plots illustrate the distribution NPC cluster g (orange) and cluster h (purple) in control (Fig. 3A) and POLG patient-derived brain organoids (Fig. 3B). In control organoids, clusters G and H appeared sparse and scattered. In contrast, POLG organoids exhibited a marked enrichment of both clusters, particularly cluster H, which formed a dense and well-defined population.

**Figure 3.**
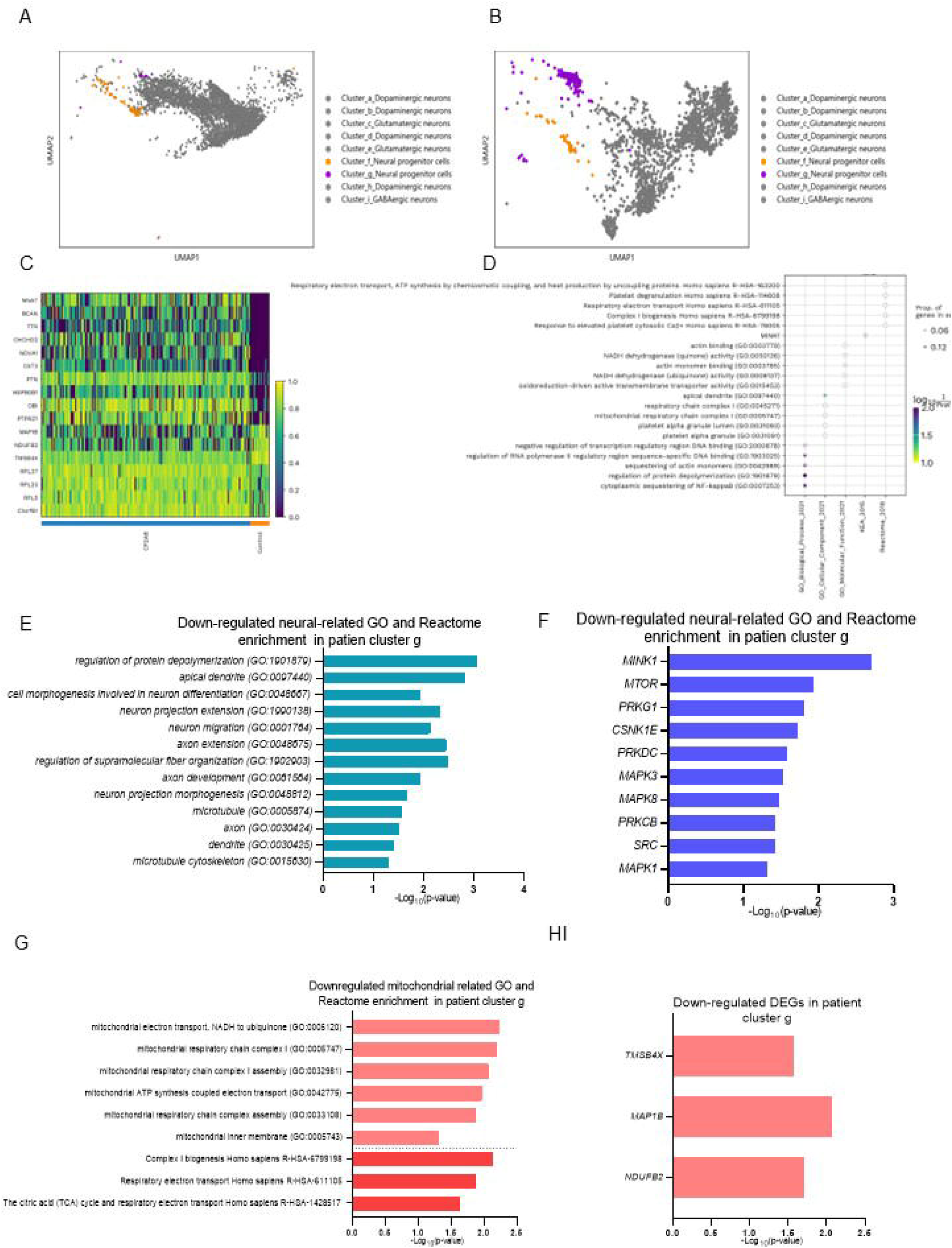
Characterization of NPC cluster g reveals transcriptional signatures of mitochondrial dysfunction and impaired neuronal development in POLG cortical organoids. (**A-B**) UMAP plots highlight the distribution of NPC subclusters g (orange) and h (purple) in control organoids (A) and POLG patient-derived organoids (B). cluster g and h are significantly enriched in POLG organoids, suggesting altered cellular states under mitochondrial stress. (**C**) Bar graph showing the proportion of NPC cluster g cells in control versus POLG organoids, with a marked increase in the patient group. (**D**) Heatmap displaying the expression of differentially expressed genes (DEGs) in cluster g between control and POLG conditions. Upregulation of stress- and inflammation-related genes such as *IFI27, CDKN1A, FOS, JUN,* and *NFKBIA* is evident in POLG NPCs. (**E**) Dot plot of GO biological processes enriched in cluster g. Upregulated terms include “response to oxidative stress,” “unfolded protein response,” and “regulation of cell death,” reflecting stress adaptation mechanisms. (**F**) Bar chart showing GO terms downregulated in POLG NPC cluster g. Suppressed functions include neuronal projection, dendrite development, and cytoskeletal organization, indicating impaired neuronal differentiation potential. (**G**) Reactome pathway analysis of downregulated genes in cluster g, highlighting suppressed pathways related to axon extension, morphogenesis, and neuron projection development. (**H**) Bar plots showing the most significantly downregulated kinases in POLG cluster g, including MINK1, MTOR, MAPK1/3/8, and PRKDC, suggesting impaired kinase signaling and synaptic function. (**I**) Additional analysis of downregulated mitochondrial-related terms, including NADH dehydrogenase complex assembly, oxidative phosphorylation, and electron transport chain components. (**G**) (J) Downregulated gene expression for *TMSB4X, MAP1B,* and *NDUFB2* in POLG NPCs, supporting the suppression of both cytoskeletal integrity and mitochondrial Complex I function.

Enrichment analysis of DEGs from NPC cluster g (Fig. S1) revealed strong associations with translational regulation, RNA processing, mitochondrial function, and neurodegenerative disease pathways. Key enriched processes included peptide chain elongation, ribosome assembly, SRP-dependent cotranslational protein targeting to the membrane, and cytoplasmic translation, indicating highly active protein synthesis machinery. Enrichment of pathways related to mRNA binding, RNA splicing (spliceosome), and nonsense-mediated decay (NMD) further suggested active RNA metabolism and quality control. Additionally, mitochondrial metabolic pathways and neurodegeneration-related pathways, such as Huntington’s disease and Parkinson’s disease signaling, were significantly enriched, pointing toward metabolic stress and increased vulnerability to neurodegenerative processes. Together, these findings suggest that NPC cluster g is characterized by elevated translational activity, robust RNA metabolism, mitochondrial dysfunction, and early signs of neurodegenerative stress.

Whereas enrichment analysis of DEGs from NPC cluster f (Fig. S2) revealed strong associations with translational regulation, RNA binding, and protein processing. Notably, pathways involved in translation initiation, cytoplasmic translation, ribosome assembly, and cotranslational protein targeting to the endoplasmic reticulum (ER) were enriched, reflecting a highly active protein synthesis system. Significant enrichment in RNA binding, mRNA binding, and rRNA binding pathways emphasized dynamic RNA processing and regulation. Furthermore, pathways related to spliceosome function, RNA transport, and nonsense-mediated decay highlighted efficient mRNA surveillance mechanisms. Enrichment of the p53 signaling pathway suggested involvement in stress responses or cell cycle regulation. Collectively, these results indicate that NPC cluster f is defined by elevated translational activity, efficient RNA metabolism, and active protein processing, supporting its role in neuronal differentiation and maintenance.

DEG analysis (Table S1) revealed distinct molecular signatures between NPC clusters g and fF. In cluster g, upregulated genes included *C1orf61, PTN, DBI, TCF12, GNG5,* and *ZFP36L1*, alongside multiple ribosomal protein genes (e.g., *RPS27, RPS19, RPL37, RPLP0, RPL13A,* and *RPS3A*), indicating elevated translational activity and potential involvement in stress response pathways. Notably, mitochondrial-related genes such as *MT-CO*1 and metabolic genes like *CKB* were also upregulated, supporting the enrichment of mitochondrial function observed in pathway analysis. In contrast, cluster f showed upregulation of genes involved in protein synthesis (e.g., F*TL, EIF1, RPS27L, RPS19*, and *RPL21*), stress response and cell cycle regulation (e.g., *DDIT3, CDKN1A, SQSTM1,* and *HSPA9*), and cytoskeletal dynamics (e.g., *VIM*). Additionally, histone-related genes such as *HIST1H1C* and *H2AFZ* were highly expressed, suggesting chromatin remodeling activities. The upregulation of multiple ribosomal proteins and RNA-binding factors further reinforces the enrichment of translation, RNA processing, and cellular stress response pathways identified in cluster f. Together, these distinct expression profiles highlight that while both clusters exhibit elevated translation-related gene expression, cluster g is more closely associated with mitochondrial function and metabolic activity, whereas cluster f reflects enhanced RNA processing, protein synthesis, and stress adaptation mechanisms, supporting their distinct biological roles during NPC development and mitochondrial dysfunction. Given these distinct transcriptional profiles, subsequent studies will prioritize further characterization of cluster g. Understanding cluster g’s specific roles in mitochondrial metabolism, translational regulation, and stress response pathways will provide critical insights into how mitochondrial dysfunction impacts neural progenitor cell fate.

Altogether, scRNA-seq analysis revealed that cortical organoid-derived NPCs form distinct clusters, with cluster g exhibiting mitochondrial and metabolic impairments, and cluster f enriched for RNA processing and stress response pathways.

### Mitochondrial dysfunction and impaired neurodevelopmental programs in POLG patient-derived NPC cluster g

Next, we compared patient organoid-derived NPC cluster g to control organoid-derived NPC cluster g. To further characterize the transcriptional features of cluster g, we performed GO and pathway enrichment analyses. The heatmap (Fig. 3C) illustrated the expression patterns of key genes across individual cells within cluster g, revealing a distinct transcriptional signature in POLG-derived cells. Notably, these cells exhibited consistent upregulation of stress- and inflammation-related genes compared to controls. Enrichment analysis of the DEGs (Fig. 3D) highlighted the top GO biological processes associated with cluster g, revealing strong enrichment for mitochondrial respiratory function, cytoskeletal organization, and transcriptional regulation. Mitochondrial-related pathways, including respiratory electron transport, ATP synthesis by chemiosmotic coupling, complex I biogenesis, and NADH dehydrogenase activity, were significantly enriched, indicating elevated mitochondrial metabolic activity. Cytoskeletal remodeling was reflected by enrichment in actin binding, actin monomer binding, and regulation of protein depolymerization, processes critical for neuronal development. Additionally, enrichment in pathways related to platelet degranulation and transcriptional regulation, including negative regulation of transcription regulatory region DNA binding and RNA polymerase II-specific DNA binding, further underscored the transcriptional reprogramming occurring within POLG-derived NPCs.

Further analysis of the downregulated DEGs in POLG patient-derived NPC cluster g (Fig. 3E, Table S2) revealed significant enrichment for pathways critical to neuronal development and cytoskeletal organization. GO terms such as regulation of protein depolymerization (GO:1901879), regulation of supramolecular fiber organization (GO:1902903), axon extension (GO:0048675), neuron projection extension (GO:1990138), neuron migration (GO:0001764), and cell morphogenesis involved in neuron differentiation (GO:0048667) were significantly suppressed, suggesting impaired neuronal morphogenesis and connectivity. Structural pathways related to axon development (GO:0061564), neuron projection morphogenesis (GO:0048812), and components of the neuronal cytoskeleton, including the microtubule network (GO:0005874), axons (GO:0030424), dendrites (GO:0030425), and the microtubule cytoskeleton (GO:0015630), were also downregulated. At the gene level (Fig. 3F, Table S3), key regulators of neuronal development and signaling, including M*INK1, MTOR, PRKG1, CSNK1E, PRKDC, MAPK3, MAPK8, PRKCB, SRC,* and *MAPK1*, were significantly downregulated. These genes are involved in critical pathways such as axonal growth, synaptic signaling, and cytoskeletal remodeling. The coordinated suppression of these genes likely contributes to impaired neuronal differentiation, projection formation, and cellular architecture maintenance in POLG NPCs.

In addition to neurodevelopmental defects, enrichment analysis of the downregulated DEGs also revealed significant suppression of mitochondrial-related pathways in POLG patient-derived NPC cluster g (Fig. 3G, Table S4). Key mitochondrial processes, including mitochondrial electron transport, NADH to ubiquinone (GO:0006120), mitochondrial respiratory chain complex I assembly (GO:0032981), and mitochondrial ATP synthesis coupled electron transport (GO:0042775), were markedly downregulated, indicating impaired oxidative phosphorylation. Structural components such as the mitochondrial respiratory chain complex I (GO:0005747) and mitochondrial inner membrane (GO:0005743) were also affected, suggesting defects in electron transport chain integrity. Reactome pathway analysis further supported these findings, with significant downregulation observed in Complex I biogenesis (R-HSA-6799198), Respiratory electron transport (R-HSA-611105), and the citric acid (TCA) cycle and respiratory electron transport (R-HSA-1428517). These pathways are essential for efficient energy production and metabolic homeostasis. The coordinated suppression of these mitochondrial pathways indicates a profound impairment of mitochondrial function in POLG NPCs, which likely exacerbates the observed neurodevelopmental deficits by limiting ATP production, increasing oxidative stress, and disrupting metabolic support for neuronal differentiation. At the gene level, key mitochondrial-related genes (Fig. 3H, Table S5), including *MAP1B, NDUFB2*, and *TMSB4X*, were significantly downregulated. MAP1B (Microtubule-associated protein 1B) is essential for axonal growth and mitochondrial trafficking, *NDUFB*2 is a core subunit of mitochondrial Complex I, critical for electron transport, and *TMSB4X* regulates cytoskeletal dynamics and is implicated in mitochondrial homeostasis. The coordinated suppression of these genes likely contributes to impaired mitochondrial function, reduced energy production, and disrupted neuronal differentiation observed in POLG NPCs. Together, these findings highlight that mitochondrial dysfunction, alongside impaired neurodevelopmental programs, underlies the vulnerability of POLG NPCs to metabolic stress and neurodegeneration.

These findings suggest POLG patient-derived NPC cluster g exhibited significant downregulation of pathways related to neuronal development, cytoskeletal organization, and mitochondrial function. Key regulators of axon growth, synaptic signaling, and mitochondrial metabolism were suppressed. These results indicate that mitochondrial dysfunction and impaired neurodevelopmental programs coexist in POLG NPC cluster g.

### Single-cell transcriptomics reveal rescue of neural progenitor imbalance and mitochondrial stress pathways

To investigate potential therapeutic strategies, we treated POLG iPSC-derived brain organoids with metformin, an established AMPK activator known to enhance mitochondrial function and cellular metabolism [6, 7]. The chemical structure of metformin is shown (Fig. 4A). Single-cell transcriptomic analysis revealed a distinct shift in cell clustering following metformin treatment in POLG organoid, as visualized by UMAP (Fig. 4B). Specifically, metformin-treated NPCs (highlighted in orange and purple) formed separate clusters from metformin treated POLG iPSC-derived brain organoids. To further investigate the effects of metformin at the cellular level, we quantified the proportion of NPC subclusters before and after treatment. In untreated POLG patient-derived cells, NPC cluster g cells were predominant (6.90%), while cluster f cells accounted for only 1.78% (Fig. 4C). Following metformin treatment, the proportion of cluster g cells was significantly reduced to 2.65%, whereas cluster f cells increased to 3.62% (Fig. 4C).

**Figure 4.**
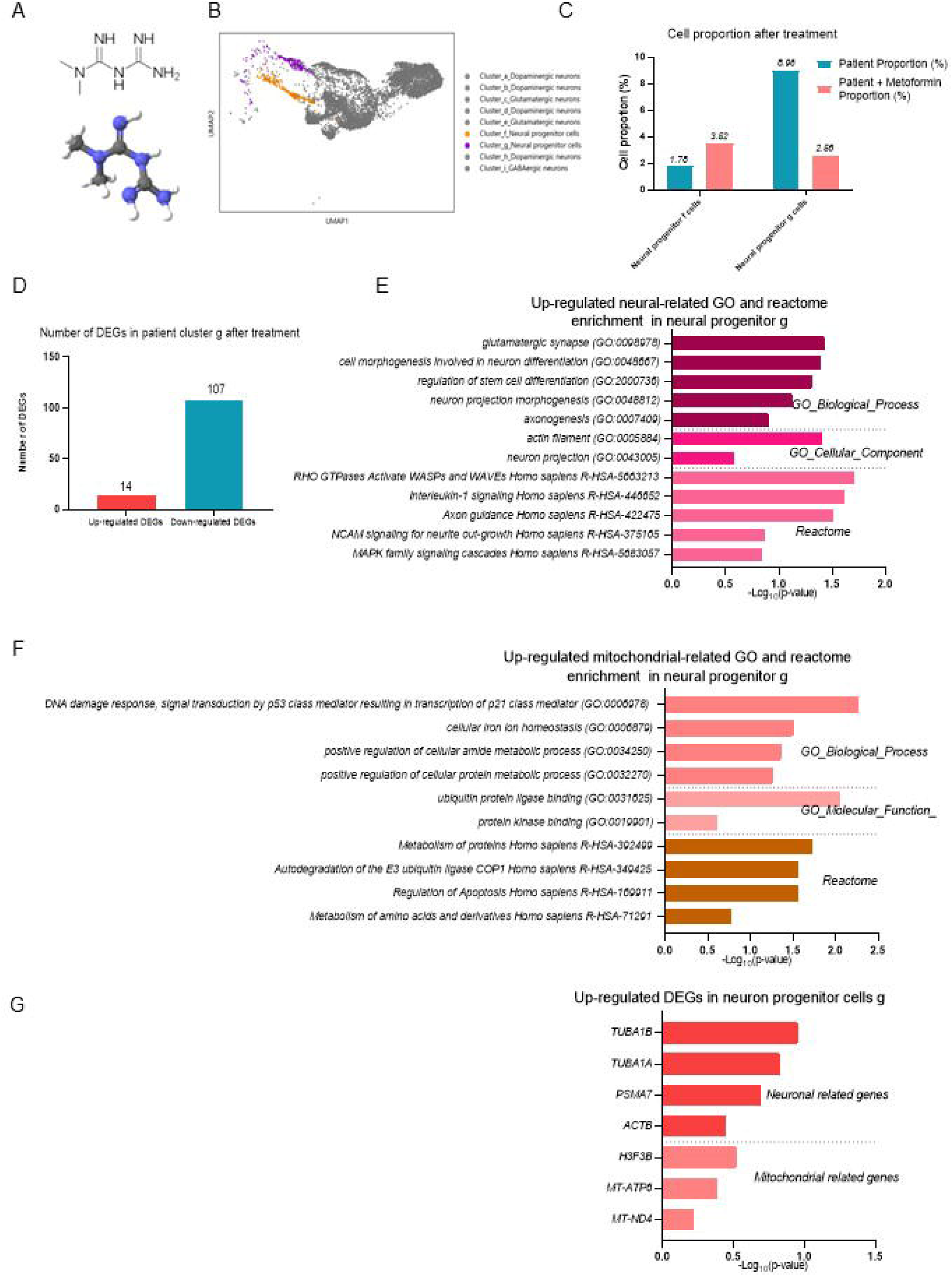
Metformin treatment reprograms neural progenitor transcriptional profiles and restores neurodevelopmental and mitochondrial pathways in POLG NPCs.. **(A)** Chemical structure of metformin, illustrating its pharmacological identity and relevance to mitochondrial modulation. (**B**) UMAP visualization of NPCs showing transcriptional reprogramming after metformin treatment. Metformin-treated NPCs (orange and purple) form distinct clusters separate from untreated POLG NPCs. (**C**) Quantification of the relative abundance of cluster g and cluster f NPCs before and after metformin treatment. Metformin reduces the proportion of pathological cluster g cells while expanding cluster f cells, suggesting a rescue of neural developmental trajectories. (**D**) **V**olcano plot of DEGs in cluster g following metformin treatment, identifying 121 DEGs (14 upregulated, 107 downregulated), indicative of broad transcriptional remodeling. (**E**) GO and Reactome pathway enrichment analysis of neural development-related genes upregulated in cluster g following metformin treatment. (**F**) GO and Reactome pathway enrichment analysis of mitochondrial function-related genes upregulated in cluster g following metformin treatment. (**G**) GO enrichment analysis of upregulated DEGs in cluster g after metformin treatment, highlighting enhanced neurodevelopmental and mitochondrial processes.

A total of 121 DEGs were identified, including 14 upregulated and 107 downregulated genes (Fig. 4D). To better understand the transcriptional response induced by metformin in neural progenitor cluster g, we performed GO and Reactome enrichment analysis of upregulated genes(Fig. 4E, Table S6). Notably, metformin treatment enhanced the expression of genes involved in key neurodevelopmental processes, including cell morphogenesis involved in neuron differentiation (GO:0048667), regulation of stem cell differentiation (GO:2000736), neuron projection morphogenesis (GO:0048812), and axonogenesis (GO:0007409). In addition, enrichment was observed in pathways related to synaptic development and cytoskeletal organization, such as glutamatergic synapse (GO:0098978), actin filament (GO:0005884), and neuron projection (GO:0043005). Reactome pathway analysis further highlighted activation of signaling cascades crucial for neuronal growth and connectivity, including RHO GTPases Activate WASPs and WAVEs (R-HSA-5663213), Interleukin-1 signaling (R-HSA-446652), Axon guidance (R-HSA-422475), NCAM signaling for neurite out-growth (R-HSA-375165), and MAPK family signaling cascades (R-HSA-5683057).

In addition to neural pathway restoration, metformin treatment also upregulated several mitochondrial-related biological processes and signaling pathways in neural progenitor cluster g (Fig. 4F, Table S7). GO enrichment analysis revealed activation of processes such as DNA damage response, signal transduction by p53 class mediator resulting in transcription of p21 class mediator (GO:0006978), cellular iron ion homeostasis (GO:0006879), positive regulation of cellular amide metabolic process (GO:0034250), and positive regulation of cellular protein metabolic process (GO:0032270). Molecular functions related to ubiquitin protein ligase binding (GO:0031625) and protein kinase binding (GO:0019901) were also significantly enriched. Reactome pathway analysis further highlighted upregulation of critical processes including Metabolism of proteins (R-HSA-392499), Regulation of Apoptosis (R-HSA-169911), Autodegradation of the E3 ubiquitin ligase COP1 (R-HSA-349425), and Metabolism of amino acids and derivatives (R-HSA-71291). Analysis of DEGs in neural progenitor cluster g after metformin treatment revealed upregulation of both neuronal- and mitochondrial-related genes (Fig. 4G, Table S8). Notably, *TUBA1B*, *TUBA1A*, *PSMA7*, and *ACTB*—genes associated with cytoskeletal structure and neuronal differentiation—were significantly upregulated, indicating enhanced neuronal development and structural organization. In addition, mitochondrial-related genes including *H3F3B*, *MT-ATP6*, and *MT-ND4* showed increased expression, reflecting improved mitochondrial function and bioenergetic capacity.

In summary, metformin treatment in POLG iPSC-derived brain organoids led to a shift in NPC subcluster distribution, reduced pathological cluster g cells, and promoted neurodevelopmental and mitochondrial pathways.

### MtDNA depletion induced by EtBr impairs NSC proliferation and neuronal differentiation

To further validate the role of mitochondrial function in neural progenitor maintenance and differentiation, we established a pharmacological model of mtDNA depletion by treating iPSC-derived neural spheres with ethidium bromide (EtBr). After 5 days of differentiation, notable morphological changes were observed. Untreated control neural spheres exhibited regular, uniform sizes with clearly defined edges (Fig. S3). In contrast, EtBr-treated spheres displayed dose-dependent alterations, including reduced sphere size, irregular boundaries, loose cellular aggregation, and increased cellular debris. These findings suggest that mitochondrial dysfunction induced by EtBr negatively impacts the viability and proliferative capacity of neural progenitor cells, emphasizing the essential role of mitochondrial integrity in maintaining normal neural differentiation and morphology.

Quantitative PCR analysis confirmed a significant, dose-dependent reduction in mtDNA copy number following EtBr treatment (Fig. S4A). Correspondingly, cell viability decreased in a dose-dependent manner (Fig. S4B), further demonstrating that EtBr severely compromises mitochondrial function, leading to impaired survival of neural progenitors.

Bright-field imaging revealed pronounced morphological changes in both NPCs and differentiated neurons upon EtBr exposure (Fig. S5A). While untreated NSCs displayed healthy proliferation with dense, compact clusters, EtBr-treated NSCs exhibited reduced proliferation, decreased cell density, disrupted morphology, and increased cellular debris. Similarly, untreated neurons formed well-defined clusters with extensive neurite outgrowth, whereas neurons from EtBr-treated conditions showed impaired differentiation, characterized by reduced neurite extension, smaller cluster size, irregular structure, and elevated cell fragmentation. These observations highlight the detrimental effects of mtDNA depletion-induced mitochondrial dysfunction on both NSC proliferation and subsequent neuronal differentiation.

Together, these findings demonstrate that mitochondrial dysfunction, whether genetically induced in POLG-derived NPC cluster g or pharmacologically induced through EtBr-mediated mtDNA depletion, leads to impaired neural progenitor proliferation, disrupted neuronal differentiation, and altered transcriptional programs. These results underscore the essential role of mitochondrial integrity in maintaining NPC identity, supporting neurodevelopmental processes, and preventing cellular stress responses.

### Metformin restores mitochondrial function without affecting neural progenitor identity in POLG NPCs

Given the cellular heterogeneity and three-dimensional complexity of cortical organoids, we then focused on a more defined and homogeneous population to precisely assess mitochondrial responses to metformin treatment. For this purpose, we utilized POLG iPSC-derived NPCs cultured in two-dimensional conditions, as previously established and characterized in our earlier studies [4, 12]. This simplified system allowed for more consistent drug exposure and facilitated quantitative assessment of mitochondrial membrane potential changes in a controlled environment. To determine the optimal concentration of metformin for mitochondrial rescue, we performed a dose-response analysis using TMRE staining to assess mitochondrial membrane potential (ΔΨm) in POLG iPSC-derived NPCs. Metformin treatment resulted in a dose-dependent increase in TMRE fluorescence intensity normalized to mitochondrial mass (measured by MitoTracker Green, MTG), with significant enhancement observed at 15.625 µM, 31.25 µM, 62.5 µM, 125 µM, and 250 µM compared to untreated controls (Fig. 5A). We further quantified total mitochondrial membrane potential across a broader concentration range using TMRE staining (Fig. 5B). A modest but statistically significant increase in TMRE intensity was detected at 125 µM and 500 µM, indicating improved mitochondrial polarization at these doses. In contrast, lower concentrations (7.8125-62.5 µM) and higher concentrations (1000-4000 µM) did not produce significant changes compared to controls. These results suggest that metformin enhances mitochondrial function in a dose-dependent manner, with optimal effects observed at intermediate concentrations. To evaluate the impact of metformin on mitochondrial mass, we performed MTG staining across the same concentration range (Fig. 5C). At lower concentrations (7.8125-125 µM), MTG fluorescence intensity remained comparable to controls. However, at higher concentrations (≥500 µM), a significant increase in mitochondrial mass was observed, with the most pronounced effect at 4000 µM.

**Figure 5.**
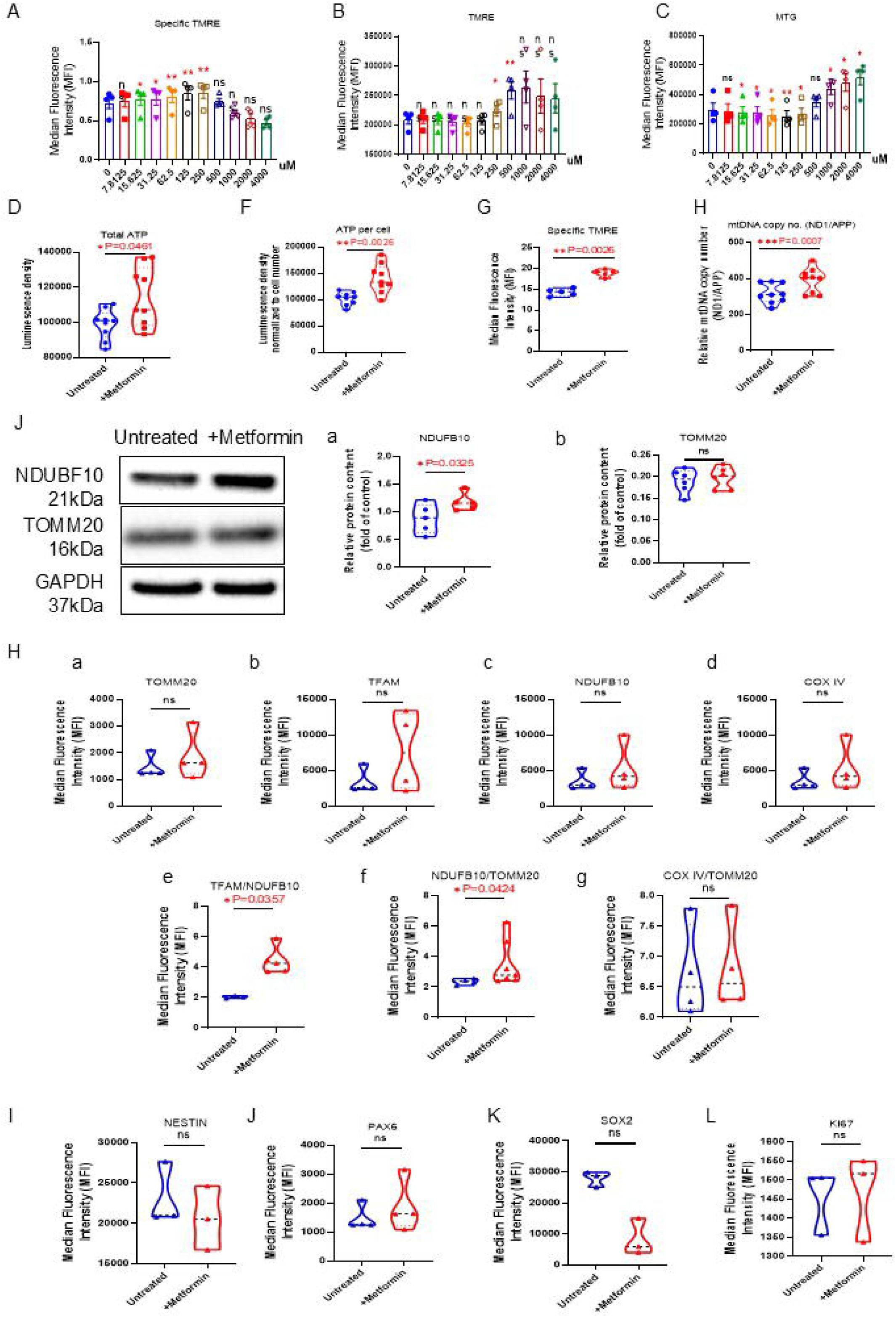
Metformin improves mitochondrial function in POLG iPSC-derived NPCs without altering progenitor identity. (**A–C**) Flow cytometry analysis revealed the dose-dependent effects of metformin on mitochondrial parameters in POLG NPCs. (A) Specific mitochondrial membrane potential per mitochondrial mass was assessed by co-staining with TMRE and MitoTracker Green (MTG). (**B**) Total mitochondrial membrane potential was evaluated by TMRE fluorescence intensity. (**C**) Mitochondrial mass was measured using MTG staining. Increasing concentrations of metformin demonstrated a selective enhancement of mitochondrial membrane potential and mass at intermediate to high doses. (**D–E**) Treatment with 250 μM metformin significantly increased (D) total intracellular ATP levels and (E) ATP content normalized per cell. (**F–G**) Quantification of mitochondrial parameters following metformin treatment: (F) Specific TMRE signal (TMRE/TOMM20) and (G) mtDNA copy number, showing enhanced mitochondrial membrane polarization and biogenesis. **(H)** Western blot analysis of mitochondrial proteins: (a) Quantification of NDUFB10 expression, and (b) Quantification of COXIV expression after metformin treatment. (**I**) Flow cytometry analysis of mitochondrial markers following metformin treatment: (a) TOMM20, (b) TFAM, (c) Specific NDUFB10, (d) COXIV, (e) Specific TMRE/TOMM20, (f) Specific NDUFB10/TOMM20, and (g) Specific COXIV/TOMM20. (**J–M**) Flow cytometry analysis of neural progenitor identity markers after metformin treatment:(J) NESTIN, (K) PAX6, (L) SOX2, and (M) proliferation marker Ki67. Mann–Whitney *U* test was used for the data presented in **D–I**. Significance is calculated by comparison to non-treated cells and is denoted for *p*-values of less than 0.05. \**p* < 0.05; \*\**p* < 0.01; \*\*\**p* < 0.001; ns, not significant.

Based on prior dose-response experiments, we selected 250 µM as the optimal concentration of metformin to treat POLG iPSC-derived NPCs. At this dose, metformin significantly improved mitochondrial function, as evidenced by a notable increase in intracellular ATP levels at total level (P = 0.0461; Fig. 5D) and specific lever per cell (P = 0.0026; Fig. 5F), indicating decreased oxidative stress. To further validate the effect of metformin on mitochondrial function and content, we assessed mitochondrial membrane potential per mitochondrial mass (specific TMRE) and mtDNA copy number in POLG iPSC-derived NPCs after treatment with 250 µM metformin. Specific TMRE signal was significantly increased (P = 0.0026, Fig. 5G), indicating that metformin enhanced mitochondrial membrane potential independently of changes in mitochondrial mass. In parallel, qPCR analysis showed a significant elevation in mtDNA copy number (P = 0.0007; Fig. 5H), suggesting that metformin also promotes mitochondrial biogenesis.

To validate the molecular effects of metformin on mitochondrial proteins, we performed Western blot analysis in POLG iPSC-derived neural progenitor cells treated with 250 µM metformin (Fig. 5J). Immunoblotting demonstrated a significant increase in NDUFB10 expression (P = 0.0325; Fig. 5J, a), a critical subunit of mitochondrial respiratory chain complex I. In contrast, the expression of TOMM20, a marker of mitochondrial mass, remained unchanged following treatment (Fig. 5J, b).

To further assess mitochondrial integrity and function in POLG iPSC-derived neural progenitor cells, we performed multiparametric flow cytometry following treatment with 250 µM metformin. Metformin did not alter the total expression levels of TOMM20 (Fig. 5H, a), a marker of mitochondrial mass, or TFAM (Fig. 5H, b), a key regulator of mitochondrial DNA transcription and replication, suggesting no major change in mitochondrial biogenesis. Similarly, total levels of NDUFB10 (Fig. 5H, c) and COXIV (Fig. 5H, d), representing respiratory chain complexes I and IV respectively, remained unchanged. However, when normalized to TOMM20, both specific TMRE (P = 0.0357; Fig. 5I, e) and specific NDUFB10 (P = 0.0424; Fig. 5H, f) were significantly increased, indicating improved mitochondrial membrane potential and respiratory complex I integrity per mitochondrial unit. In contrast, specific COXIV (Fig. 5H, g) did not show a significant change.

To determine whether metformin treatment alters the neural progenitor identity of POLG iPSC-derived NPCs, we assessed the expression of key neural markers by flow cytometry (Fig. 5I-L). NESTIN expression (Fig. 5I), a canonical neural progenitor marker, showed no significant difference between untreated and metformin-treated cells, indicating that the progenitor state was preserved. Similarly, SOX2 (Fig. 5J), a marker of stemness and early progenitor status, remained unchanged. PAX6 expression (Fig. 5K), associated with neuroectodermal commitment, also did not differ between groups, suggesting no shift in lineage specification. Additionally, Ki67 levels (Fig. 5L), a marker of cell proliferation, were unaltered, indicating that metformin did not affect proliferative capacity.

Together, treatment with 250 µM metformin significantly improved mitochondrial function in POLG iPSC-derived NPCs, as evidenced by enhanced mitochondrial membrane potential, increased ATP production, elevated mtDNA copy number, and improved respiratory chain integrity. Importantly, metformin treatment did not alter neural progenitor identity or proliferative capacity, indicating selective restoration of mitochondrial activity without affecting cellular differentiation status.

### Metformin enhances early neural induction and sustains mitochondrial function during neuronal differentiation in POLG iPSC-derived cells

To determine whether metformin influences early neural differentiation, we treated human iPSCs with 250 μM metformin during the initial five days of neural induction and assessed neural aggregate morphology and growth (Fig. 6A-C). Bright-field imaging revealed that metformin-treated cultures developed larger and more compact neural rosettes compared to untreated controls (Fig. 6A). Quantification confirmed a significant increase in both aggregate diameter (P = 0.0497; Fig. 6B) and total area (P = 0.0054; Fig. 6C), indicating enhanced neural progenitor expansion and improved efficiency of neural induction.

**Figure 6.**
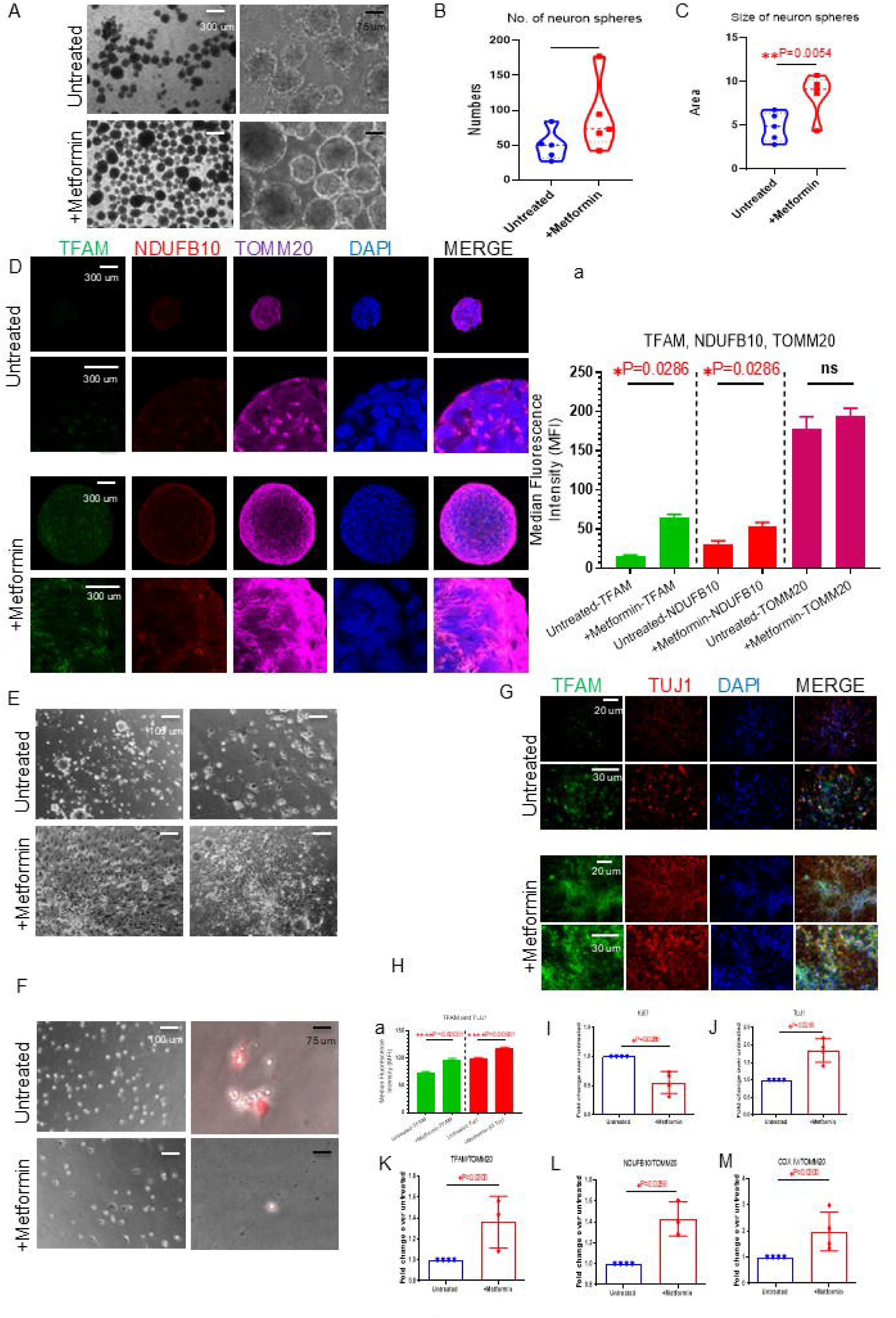
Metformin promotes early neural induction and supports mitochondrial function during neuronal differentiation in POLG iPSC-derived cultures. (**A**) Bright-field images of neural aggregates at Day 5 of induction from untreated (top) and metformin-treated (bottom) cultures. Metformin-treated aggregates appeared more compact and larger. Scale bars: 300 μm (white) and 75 μm (black). (**B–C**) Quantification of neural aggregate morphology at Day 5, showing (B) number of aggregates and (C) aggregate size. Metformin treatment significantly enhanced early neuroectodermal induction. (**D**) Representative immunofluorescence images of neural aggregates stained for TFAM (green), NDUFB10 (red), TOMM20 (magenta), and DAPI (blue). Metformin treatment increased mitochondrial protein expression. Scale bars: 300 μm. (a) Quantification of TFAM, NDUFB10, and TOMM20 fluorescence intensity. (**E**) Bright-field images of differentiated neurons derived from untreated and metformin-treated neural aggregates. Scale bars: 100 μm. (**F**) Bright-field and Ethidium Homodimer-1 (EthD-1) staining of adherent cultures showing reduced dead cell signals in metformin-treated neurons. Scale bars: 300 μm (white) and 75 μm (black). (**G–H**) (G) Immunostaining and (H) quantification of differentiated neurons for neuronal marker TUJ1 (red) and mitochondrial marker TFAM (green). DAPI (blue) marks nuclei. Metformin-treated neurons showed increased TFAM signal along neuronal processes. Scale bars: 20–30 μm. (**I–M**) Flow cytometry analysis of differentiated neurons comparing metformin-treated and untreated POLG neurons, showing expression of: (I) Ki67, (J) TUJ1, (K) TFAM/TOMM20 ratio, (L) NDUFB10/TOMM20 ratio, and (M) COXIV/TOMM20 ratio. Mann–Whitney *U* test was used for the data presented in **B, C, D, F–M**. Significance is calculated by comparison to non-treated cells and is denoted for *p*-values of less than 0.05. \**p* < 0.05; \*\**p* < 0.01; \*\*\*\**p* < 0.000; ns, not significant.

To examine the effects of metformin on mitochondrial function during early neurodevelopment, we performed immunocytochemistry for mitochondrial markers in day 5 neural aggregates (Fig. 6D). Compared to controls, metformin-treated aggregates exhibited significantly increased expression of TFAM (green), a key regulator of mitochondrial DNA maintenance, and NDUFB10 (red), a complex I subunit. In contrast, TOMM20 (magenta), a marker of mitochondrial mass, remained unchanged between groups (Fig. 6D, a). The quantification confirmed that TFAM and NDUFB10 fluorescence intensities were significantly elevated (P = 0.0286 for both; Fig. 6D, a), suggesting that metformin enhances mitochondrial gene expression and respiratory chain integrity during early neural fate acquisition, independent of changes in mitochondrial abundance.

To assess the functional consequences of early metformin treatment, neural aggregates were plated for adherent differentiation (Fig. 6E). While both groups adhered and extended neurites over time, metformin-treated aggregates showed enhanced attachment and more robust neurite outgrowth. Ethidium Homodimer I (EthD-I) staining (Fig. 6F) revealed a reduced proportion of dead or membrane-compromised cells in metformin-treated cultures, suggesting improved viability during neural differentiation. These findings indicate that early metformin exposure promotes not only neural induction and mitochondrial function but also supports neuronal survival and differentiation efficiency.

We next evaluated the persistence of these effects during terminal differentiation. Immunocytochemistry for the neuronal marker TUJ1 (Fig. 6G) demonstrated strong neuronal identity in both groups, with visibly increased TFAM signal in metformin-treated neurons (P = 0.00001, Fig. 6G&H). Merged images showed widespread TFAM localization along neuronal processes. Quantification of TFAM fluorescence revealed significantly elevated levels in metformin-treated NPCs and an even greater increase in TUJ1⁺ neurons (P = 0.00001; Fig. 6G&H), confirming that metformin enhances mitochondrial transcriptional activity throughout differentiation.

To determine whether metformin influences cell proliferation at the protein level, we assessed Ki67 expression by flow cytometry in POLG iPSC-differentiated neurons. Although metformin-treated neurons exhibited a downward trend in Ki67 expression, the difference was statistically significant (P = 0.0286, Fig. 6I), suggesting that metformin does not markedly alter proliferative status during the neural progenitor stage. To further evaluate mitochondrial quality in differentiated neurons, we performed flow cytometry analysis for TUJ1, and mitochondrial markers normalized to TOMM20. Metformin-treated neurons showed a significant increase in TUJ1 expression (P = 0.0286, Fig. 6J), confirming enhanced neuronal identity. Additionally, the ratios of TFAM/TOMM20 (P = 0.030, Fig. 6K), NDUFB10/TOMM20 (P = 0.0286, Fig. 6L), and COXIV/TOMM20 (P = 0.030, Fig. 6M) were all significantly elevated, indicating improved mitochondrial transcriptional activity and restoration of respiratory chain complex I and IV per mitochondrial unit.

Early metformin treatment during neural induction enhanced neural progenitor expansion, improved mitochondrial gene expression and respiratory chain integrity, and promoted neuronal survival and differentiation efficiency. These effects persisted through terminal differentiation, with metformin-treated neurons exhibiting elevated mitochondrial transcriptional activity, enhanced neuronal identity, and restored mitochondrial respiratory complex function, without markedly altering proliferative status.

### Metformin promotes mitochondrial biogenesis, fusion, and quality control in POLG NPCs via mTOR inhibition, AMPK activation, and enhanced mitophagy

To further elucidate the molecular mechanisms underlying metformin-induced mitochondrial improvements, we next investigated key signaling pathways involved in mitochondrial biogenesis, dynamics, and quality control in POLG iPSC-derived NPCs. To investigate the upstream regulatory mechanisms by which metformin promotes mitochondrial biogenesis, we assessed the expression of key signaling molecules involved in metabolic and mitochondrial regulation, including phosphorylated mTOR (p-mTOR), phosphorylated SIRT1 (p-SIRT1), SIRT3, phosphorylated AMPK (p-AMPK), NFR1, and PGC-1α in POLG iPSC-derived NPCs (Fig. 7A). Metformin treatment significantly reduced the p-mTOR/mTOR ratio (P = 0.0014; Fig. 7A, a), indicating effective inhibition of the mTOR signaling pathway. In contrast, phosphorylation levels of SIRT1 remained unchanged (Fig. 7A, b), suggesting that SIRT1 is not directly involved in the metformin-mediated response in this context. Importantly, PGC-1α-a central transcriptional coactivator regulating mitochondrial biogenesis-was significantly upregulated following metformin treatment (P = 0.0094; Fig. 7A, c). Metformin treatment significantly increased the expression of NRF1, a key transcription factor involved in mitochondrial biogenesis, in POLG iPSC-derived NPCs (P = 0.015; Fig. 7A, d). In addition, both SIRT3 protein level (P = 0.0129; Fig. 7A, e) and the p-AMPK/AMPK ratio (P = 0.0159; Fig. 7A, f) were elevated, indicating activation of mitochondrial regulatory programs and energy-sensing pathways.

**Figure 7.**
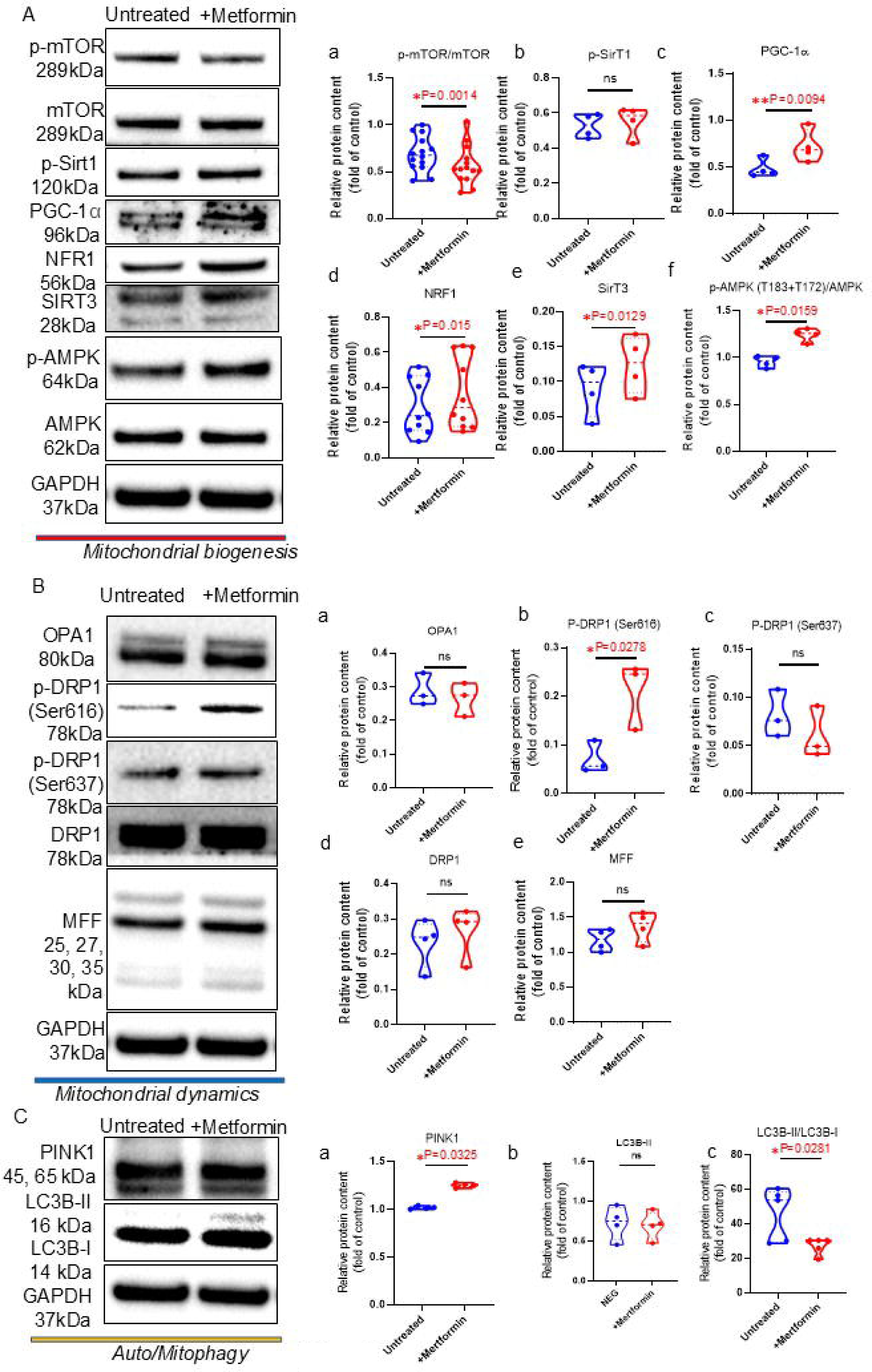
Metformin activates mitochondrial biogenesis, fusion, and mitophagy pathways in POLG NPCs via mTOR inhibition and AMPK/SIRT3 activation. **(A)** Western blot analysis of downstream signaling pathways regulating mitochondrial biogenesis in POLG iPSC-derived neural progenitor cells. Metformin treatment reduced p-mTOR/mTOR ratio, while increasing the expression of NRF1 and PGC-1α, and elevating SIRT3 levels and p-AMPK/AMPK ratio. Phosphorylation of SIRT1 remained unchanged. **(B)** Western blot analysis of mitochondrial dynamics-related proteins. Metformin increased phosphorylation of DRP1 at Ser637, a modification that inhibits mitochondrial fission, without altering total DRP1, OPA1, or MFF expression. Phosphorylation of DRP1 at Ser616, which promotes fission, remained unchanged. **(C)** Western blot and quantification of mitophagy markers. Metformin significantly upregulated PINK1 and increased the LC3B-II/LC3B-I ratio, indicative of enhanced mitochondrial quality control through mitophagy. Total LC3B levels were not significantly affected. Mann–Whitney *U* test was used for the data presented in **B, C, D, F–M**. Significance is calculated by comparison to non-treated cells and is denoted for *p*-values of less than 0.05. \**p* < 0.05; \*\**p* < 0.01; ns, not significant.

To further assess the effects of metformin on mitochondrial dynamics, we examined the expression of fusion and fission regulators, including OPA1, p-DRP1 (Ser616), and p-DRP1 (Ser637) (P = 0.0278; Fig. 7B). While OPA1 levels exhibited a non-significant upward trend (Fig. 7B, a), phosphorylation of DRP1 at Ser637 was significantly increased (P = 0.0278; Fig. 7B, b), a modification known to inhibit DRP1 activity and reduce mitochondrial fission. In contrast, levels of p-DRP1 (Ser616), which promotes mitochondrial fragmentation, remained unchanged (Fig. 7B, c). Additionally, total levels of DRP1 and its adaptor MFF (Fig. 7B, d-e) were unaffected by metformin, indicating that metformin modulates mitochondrial dynamics via post-translational regulation of DRP1 activity rather than through changes in overall protein abundance.

To explore the role of metformin in mitochondrial quality control and autophagy, we next assessed PINK1 and LC3B expression (Fig. 7C). PINK1, a critical initiator of mitophagy, was significantly upregulated in metformin-treated NPCs (P = 0.0325; Fig. 7C, a), suggesting enhanced mitochondrial surveillance. Moreover, metformin increased the expression of LC3B-II (Fig. 7C, b), the lipidated form associated with autophagosome formation. Although total LC3B levels remained unchanged, the LC3B-II/LC3B-I ratio was significantly elevated (P = 0.0281; Fig. 7C, c), an indicative of increased autophagic flux. Together, these results demonstrate that metformin activates PINK1-mediated mitophagy and promotes autophagy-dependent mitochondrial clearance, thereby contributing to enhanced mitochondrial homeostasis in POLG iPSC-derived neural progenitors.

Together, these findings demonstrate that metformin enhances mitochondrial homeostasis in POLG iPSC-derived NPCs through coordinated activation of AMPK–PGC-1α–SIRT3 signaling, inhibition of mTOR, promotion of mitochondrial fusion, and stimulation of PINK1-mediated mitophagy. By targeting multiple aspects of mitochondrial regulation, metformin restores mitochondrial integrity and quality control mechanisms critical for neural progenitor health.

## Discussion

In this study, we established a robust 3D cortical organoid model from POLG patient-derived iPSCs carrying compound heterozygous mutations (A467T and W748S), enabling the recapitulation of early human cortical development and the modeling of mitochondrial neurodevelopmental pathology. These organoids exhibited well-structured cortical architecture, expressing key layer-specific neuronal and glial markers, and revealed developmental abnormalities consistent with POLG-related disorders, including delayed differentiation and altered progenitor dynamics.

Single-cell transcriptomic profiling provided critical insights into the cellular heterogeneity within POLG organoids. We identified two distinct NPC clusters: cluster f, associated with neuronal differentiation, and cluster g, enriched in stress- and inflammation-related genes such as *IFI27, CDKN1A, FOS, JUN,* and *NFKBIA*. POLG organoids showed a marked expansion of cluster g and an overall increase in NPC abundance compared to controls, indicating a differentiation block. Gene ontology and Reactome analyses revealed widespread downregulation of mitochondrial pathways—particularly those involved in oxidative phosphorylation, electron transport, and complex I assembly (e.g., NDUFB2, MT-ND4)—as well as suppression of neurodevelopmental processes such as axonogenesis, neuronal projection, and cytoskeletal remodeling. These findings support the hypothesis that POLG mutations impair neural differentiation by inducing mitochondrial and metabolic stress that shifts NPCs toward a dysfunctional, pro-inflammatory phenotype, echoing prior findings in mitochondrial diseases and neurodevelopmental disorders [1, 13, 14].

Metformin treatment resulted in a dramatic shift in NPC subpopulation dynamics and transcriptional programming. The proportion of stress-associated cluster g NPCs was reduced by more than half (from 6.9% to 2.65%), while cluster f expanded (from 1.78% to 3.62%), suggesting a partial restoration of neurogenic capacity. Transcriptomic analyses showed that metformin reactivated suppressed neurodevelopmental pathways, including neuronal projection, synapse formation, and cytoskeletal organization. Furthermore, metformin restores key signaling networks—such as MAPK, NCAM, and Rho GTPase pathways—that are essential for neurite extension and synaptic connectivity. These results are consistent with prior studies showing that metformin enhances neurogenesis in models of aging and injury [15, 16], and they extend this effect to a human genetic model of mitochondrial dysfunction.

In parallel, metformin significantly improved mitochondrial function in POLG NPCs. Functional assays demonstrated a dose-dependent increase in mitochondrial membrane potential (ΔΨm), with optimal effects at 250 µM, accompanied by elevated ATP levels. Notably, these improvements occurred without changes in mitochondrial mass (TOMM20), but were associated with increased mtDNA copy number and enhanced expression of respiratory chain subunits NDUFB10 and COXIV, indicating improved mitochondrial quality per organelle. Mechanistically, metformin promoted mitochondrial biogenesis through coordinated inhibition of mTOR signaling and activation of AMPK and PGC-1α [17, 18]. Interestingly, SIRT1 phosphorylation remained unchanged, but SIRT3, a mitochondrial-localized deacetylase, was significantly upregulated. SIRT3 is known to enhance mitochondrial respiration and antioxidant defense via deacetylation of mitochondrial enzymes [19], and its elevation suggests a key role in metformin-mediated mitochondrial rescue in POLG NPCs.

Mitochondrial dynamics—the balance between fission and fusion—is critical for maintaining mitochondrial integrity, energy production, and neuronal health [20]. Dysregulation of these processes is commonly observed in various neurodegenerative and mitochondrial diseases [21]. In this study, we found that metformin treatment significantly increased phosphorylation of dynamin-related protein 1 (DRP1) at Ser637, a well-known inhibitory site that reduces DRP1-mediated mitochondrial fission. This post-translational modification favors mitochondrial elongation and fusion without altering total DRP1 or its adaptor protein MFF levels. Enhanced phosphorylation of DRP1 at Ser637 suggests that metformin suppresses mitochondrial fragmentation, a pathological feature observed in models of Alzheimer’s disease and mitochondrial disorders [21, 22].

Excessive mitochondrial fission leads to the production of fragmented, dysfunctional mitochondria, which are less efficient in ATP generation and more prone to generating reactive oxygen species (ROS). By promoting mitochondrial fusion, metformin likely preserves mitochondrial function, maintains bioenergetic homeostasis, and prevents the accumulation of defective organelles. Promoting a fused mitochondrial network also improves mitochondrial complementation, metabolic flexibility, and supports neuronal survival under stress conditions.

Finally, we show that metformin enhances mitochondrial quality control through upregulation of PINK1 and increased LC3B-II/LC3B-I ratio, indicating activation of mitophagy and autophagic flux [23, 24]. These changes suggest that metformin supports mitochondrial health not only by promoting biogenesis and fusion but also by facilitating the removal of damaged mitochondria.

Compared to other pharmacological strategies, metformin provides a clinically accessible, multi-targeted approach that simultaneously enhances mitochondrial function and restores developmental programs, supporting its therapeutic potential in mitochondrial neurodevelopmental disorders.

## Declarations Ethics approval

The project was approved by the Western Norway Committee for Ethics in Health Research (REK nr. 2012/919).

## Availability of data and materials

The datasets generated and analyzed during the study are included with Supplemental Information. The RNA sequencing analysis read count data can be accessed in NCBI Gene Expression Omnibus (GEO) data deposit system with an accession number GEO GSE234659. All other data is available from the corresponding author upon request.

## Author’s contributions

K.L contribute to the conceptualization; A.C contribute to the methodology; A.C, S.E.S, C.K.K contribute to the investigation; K.L and A.C contribute to the writing original draft; all authors contribute to writing review and editing; K.L contribute to the funding acquisition and to the resources; K.L contributes to the supervision. All authors agree with the authorships.

## Acknowledgements

We thank members of the Molecular Imaging Centre and Flow Cytometry Core Facility for their expertise and assistance in confocal imaging and flow cytometry data recording.

---

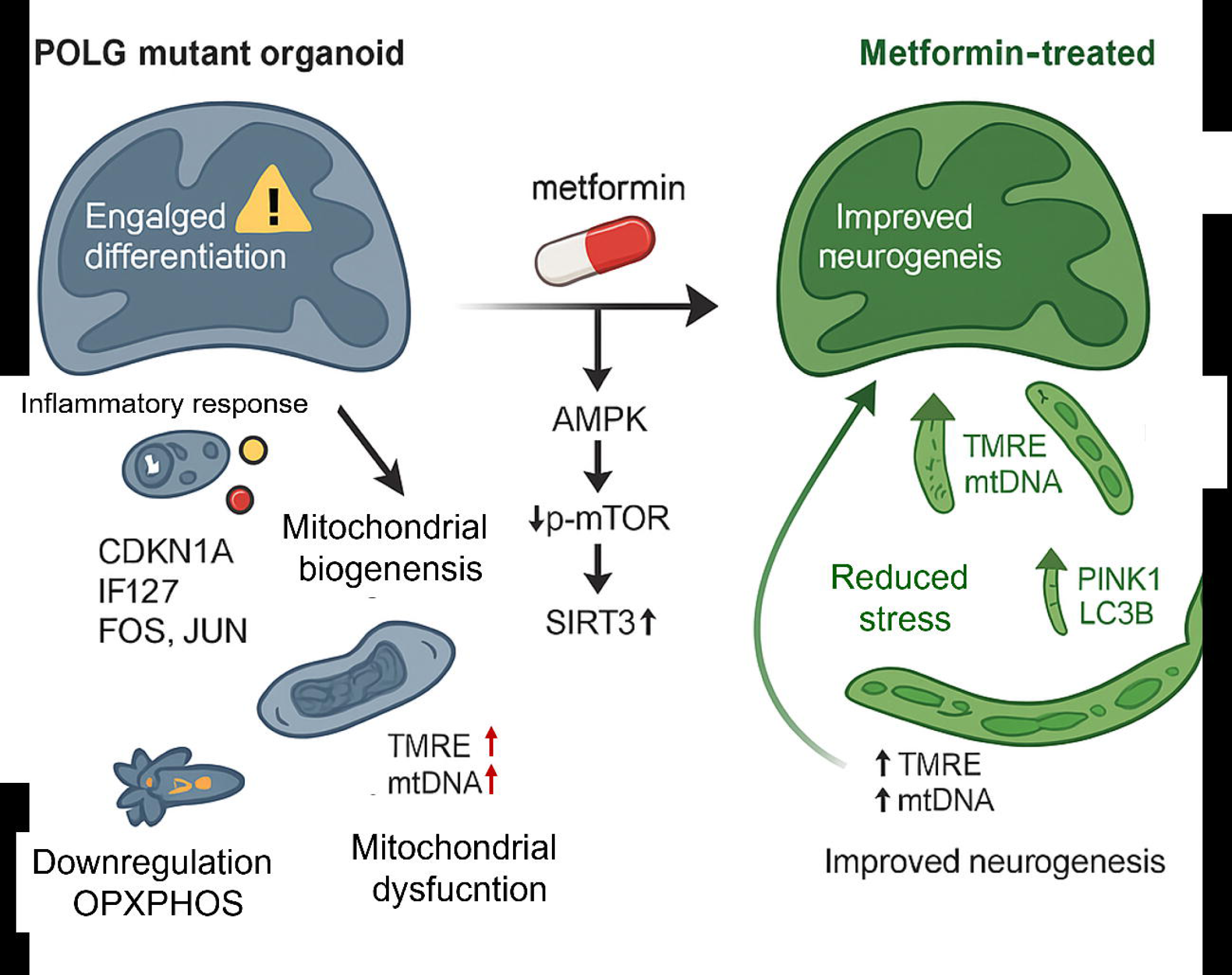

