## Supplemental file for "Metformin boosts mitochondria and neurogenesis via AMPK/mTOR/SIRT3 in POLG mutant organoids"

### Supplemental Figures

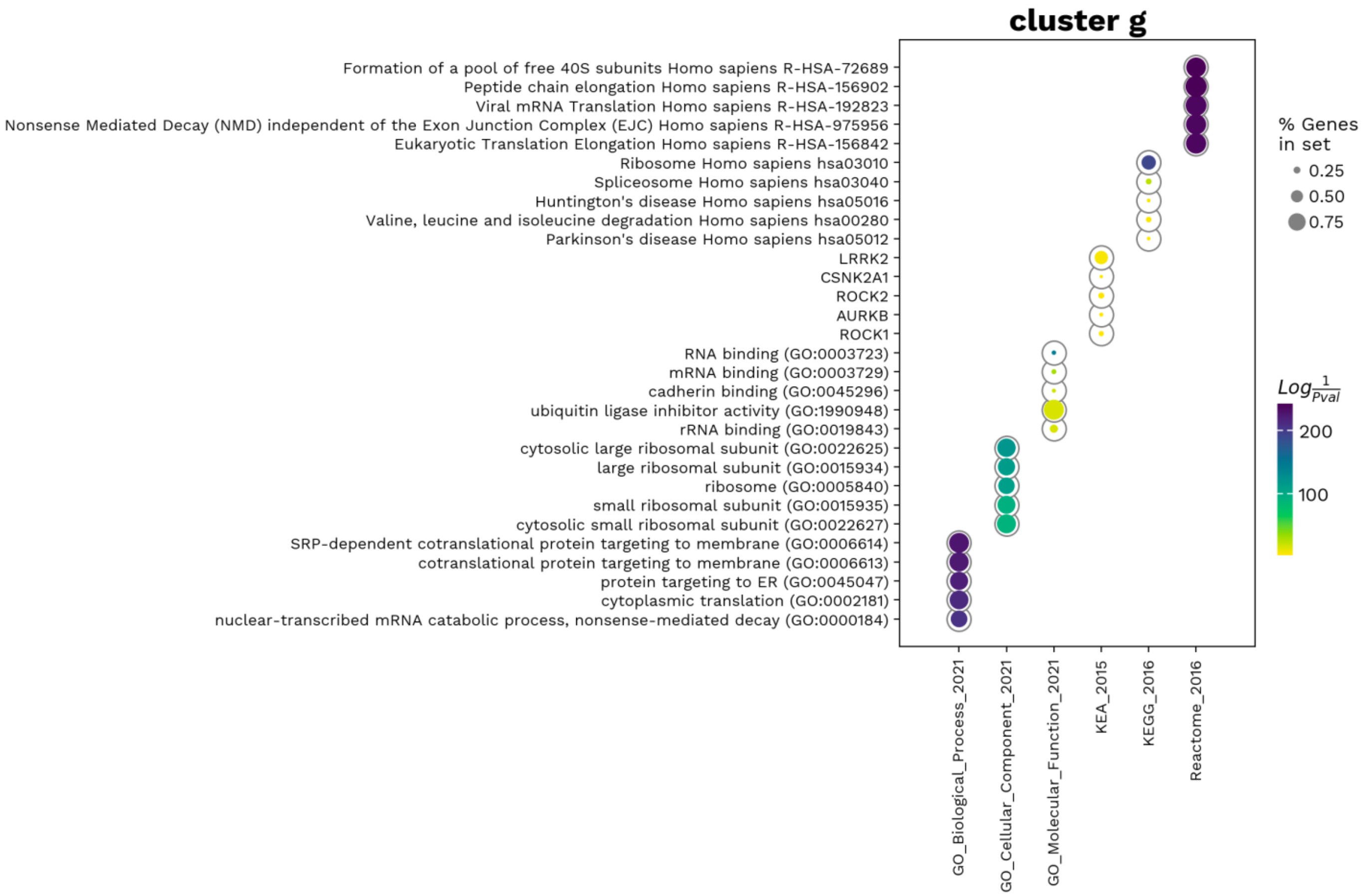


**Figure S1. Functional enrichment analysis of transcription-factor-related activities in cluster g.**

UMAP scatter plot displaying significantly enriched GO terms related to transcription factor activity. Dot size represents the proportion of genes enriched in each GO term, and color intensity (from purple to yellow) corresponds to enrichment strength.

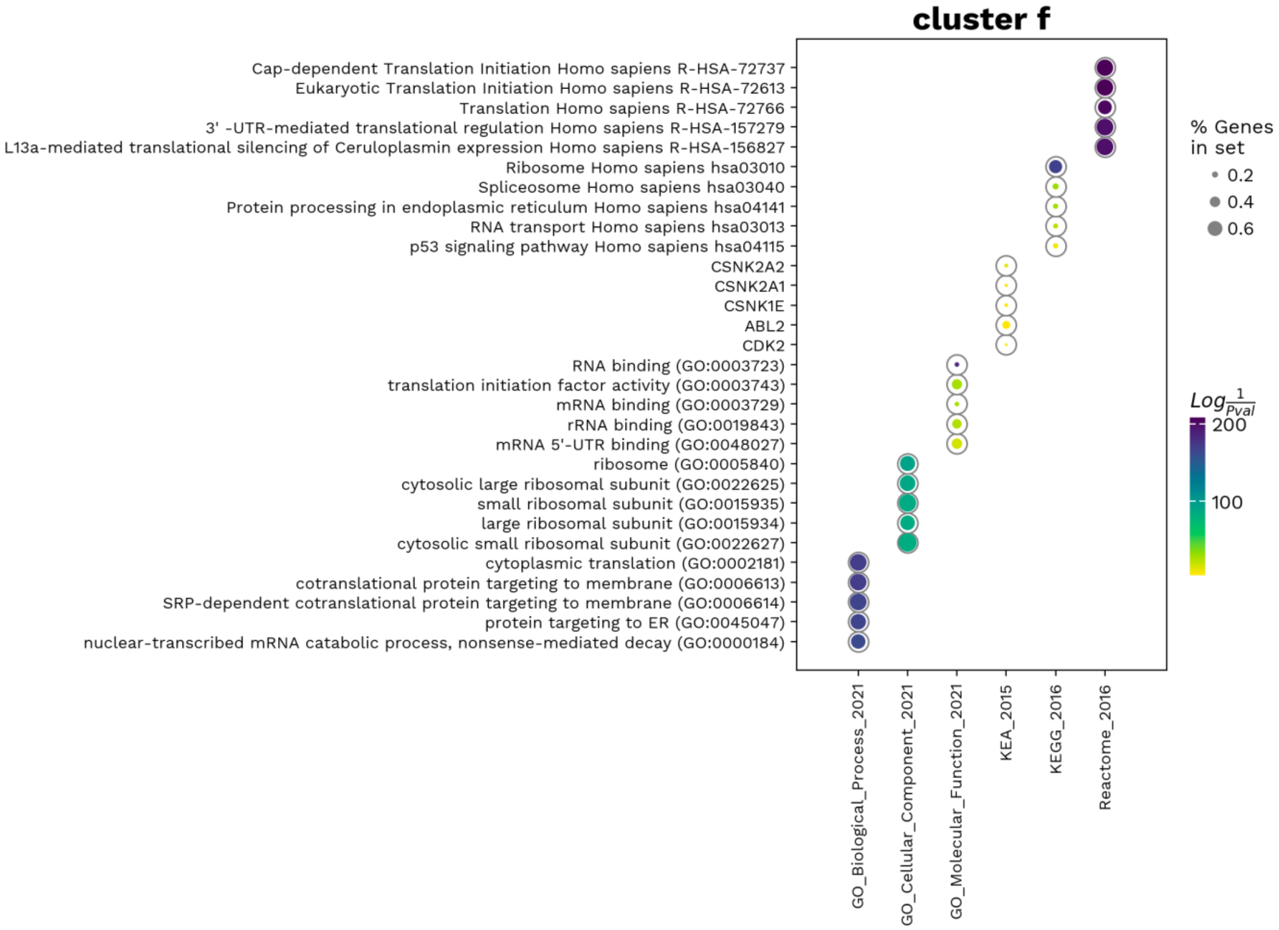


**Figure S2. Functional enrichment analysis of transcription-factor-related activities in cluster f.**

UMAP scatter plot displaying significantly enriched GO terms related to transcription factor activity. Dot size represents the proportion of genes enriched in each GO term, and color intensity (from purple to yellow) corresponds to enrichment strength.

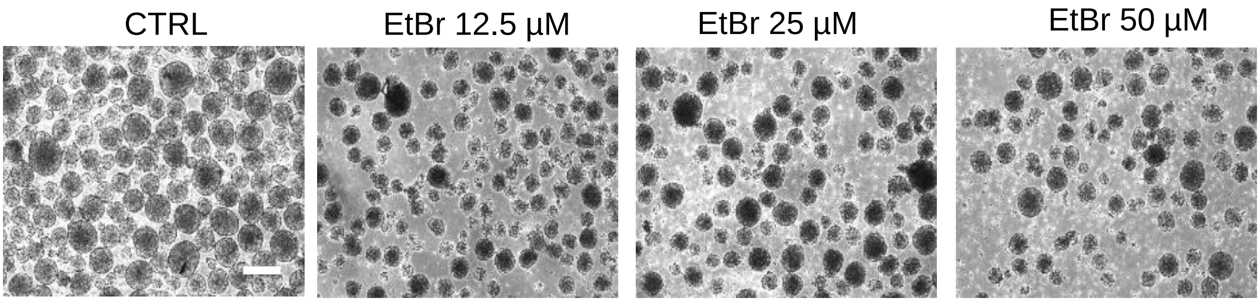


**Figure S3. Morphological changes of neural spheres derived from control iPSCs following EtBr treatment.**

Representative bright-field images showing neural spheres differentiated from control iPSCs at day 5, treated with ethidium bromide (EtBr). Panels represent increasing concentrations of EtBr treatment (A: Control untreated; B-D: increasing EtBr concentrations). Morphological observations indicate dose-dependent alterations, including decreased sphere size, irregular shapes, and increased cellular fragmentation. Scale bar: 100 µm.

A

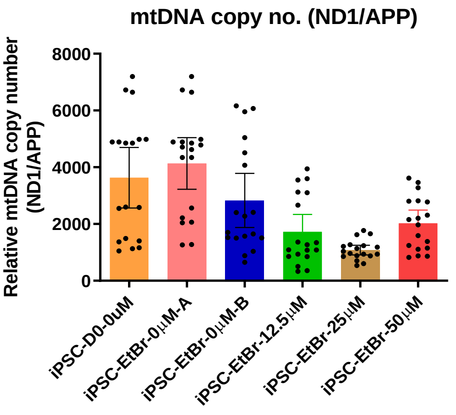


B

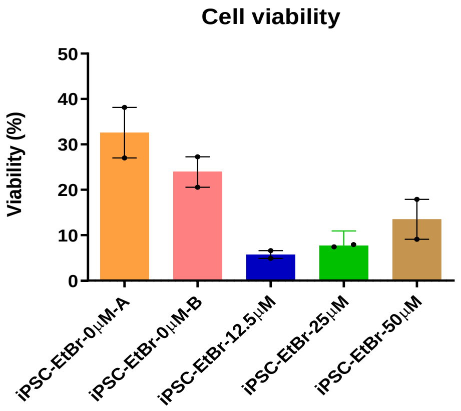


**Figure S4. mtDNA content and cellular viability of neural spheres following EtBr treatment.**

(A) Quantitative PCR analysis demonstrating significant, dose-dependent reduction of mitochondrial DNA (mtDNA) copy number after EtBr exposure. (B) Cell viability measured by trypan blue staining. Data represent mean ± SD, with individual replicates indicated.

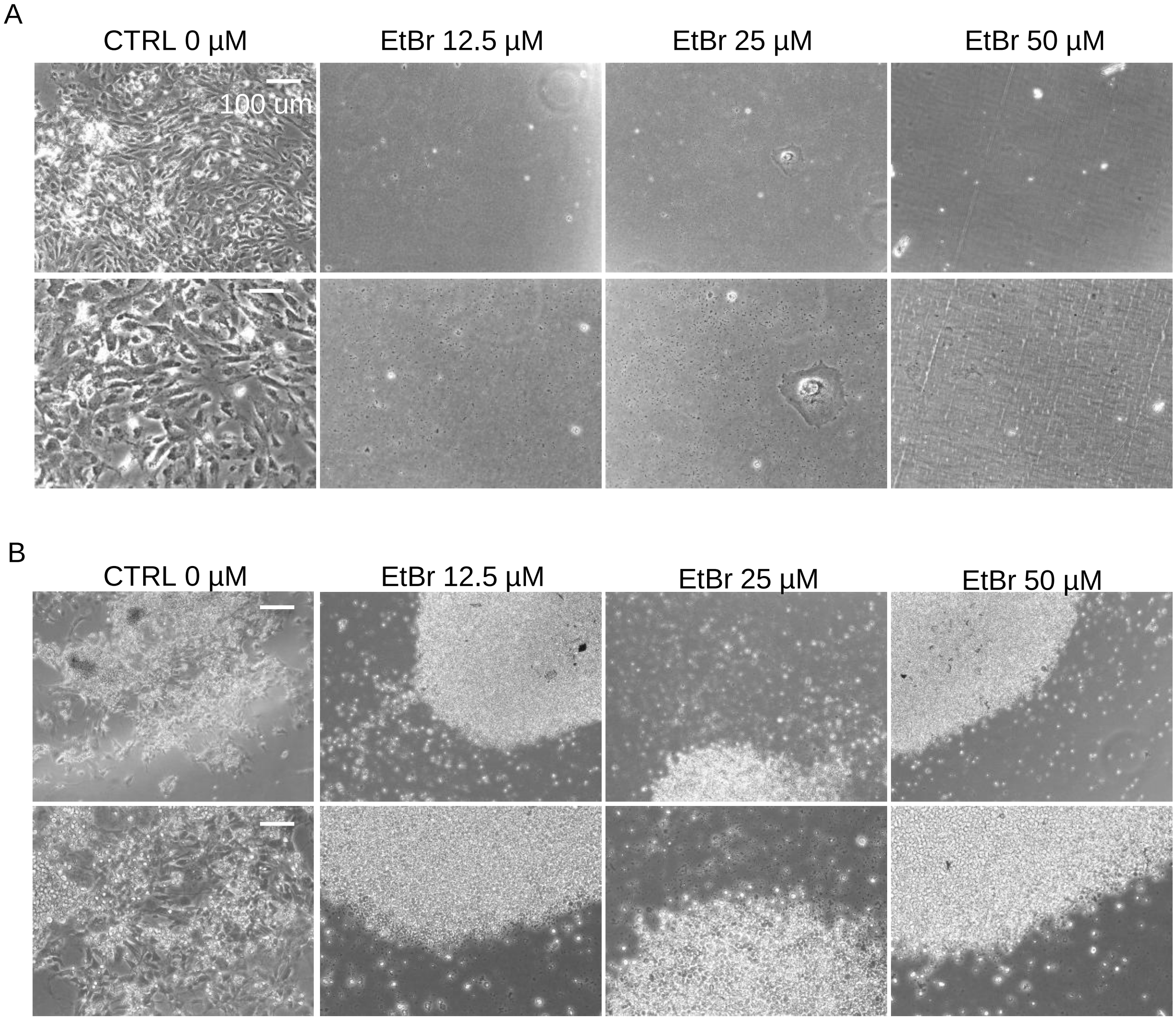


**Figure S5. Morphological effects of EtBr treatment on neural stem cell (NSC) proliferation and neuronal differentiation.**

Representative bright-field images illustrating the impact of increasing EtBr concentrations on NPCs (top panels) and differentiated neurons (bottom panels). Untreated control cells exhibited healthy proliferation (NPCs) and robust neuronal differentiation. Scale bar: 100 µm.

Supplemental Tables

**Table S1. DEGs in neuron progenitor g and f.**

| NPC g | | NPC f | |
|---|---|---|---|
| Genes | logfoldchanges | Genes | logfoldchanges |
| *C1orf61* | 4.32780695 | *FTL* | 2.63744998 |
| *PTN* | 3.822497606 | *EIF1* | 1.941274524 |
| *DBI* | 3.581779242 | *DDIT3* | 4.387285233 |
| *RPS27* | 1.586322546 | *CDKN1A* | 4.384426117 |
| *RPS19* | 1.425219417 | *RPS27L* | 2.436941147 |
| *RPL37* | 1.705489159 | *H3F3B* | 1.35811615 |
| *EEF1A1* | 1.311109185 | *SLC3A2* | 3.059803724 |
| *RPLP0* | 1.717797995 | *TRMT112* | 1.898112893 |
| *TCF12* | 4.008550644 | *FTH1* | 1.265012026 |
| *RPS27L* | 2.761666536 | *RPS19* | 1.016398311 |
| *GNG5* | 3.4569242 | *RPS27* | 1.091330886 |
| *RPL13A* | 1.498371124 | *RPL21* | 1.08894825 |
| *RPL13* | 1.436141253 | *SQSTM1* | 3.185659885 |
| *RPS3A* | 1.319384933 | *RPS13* | 0.98312217 |
| *RPL27A* | 1.369380593 | *VIM* | 2.68647933 |
| *GPM6B* | 2.651038408 | *HIST1H1C* | 4.467734814 |
| *RPLP1* | 1.162662625 | *RPL41* | 1.148336053 |
| *RPL7* | 1.616607308 | *B2M* | 2.880173445 |
| *RPS6* | 1.782844305 | *ZFAS1* | 1.67312336 |
| *RPL5* | 1.492417097 | *RPL18A* | 0.866674006 |
| *RPS2* | 1.118875265 | *CLU* | 1.924841523 |
| *PTPRZ1* | 3.151057005 | *HSPA9* | 1.870765686 |
| *LINC01158* | 3.268900394 | *H2AFZ* | 1.139682174 |
| *RPL18A* | 1.268128276 | *EIF5* | 1.367935538 |
| *MT-CO1* | 1.173237801 | *RPLP1* | 0.68926543 |
| *CKB* | 1.926262617 | *RPL27A* | 0.850938797 |
| *RPS14* | 1.171328545 | *HSPE1* | 1.121474862 |
| *RPL3* | 1.56616962 | *C6orf48* | 1.705809832 |
| *RPL35A* | 1.304636598 | *HERPUD1* | 2.49187398 |
| *ZFP36L1* | 4.616969585 | *GAS5* | 1.935317039 |

**Table S2. Mitochondrial related GO enriched pathways downregulated in neuron progenitor g neuron cluster, related to Fig. 3E.**

| Gene_set | Term | P-value | Adjusted p-value | Logfoldchanges |
|---|---|---|---|---|
| Go_biological_process_2021 | Regulation of protein depolymerization (GO:1901879) | 0.000899754 | 0.010330529 | 3.045876272 |
| Go_biological_process_2021 | Regulation of supramolecular fiber organization (GO:1902903) | 0.003446153 | 0.011039208 | 2.462665438 |
| Go_biological_process_2021 | Axon extension (GO:0048675) | 0.003595807 | 0.011039208 | 2.444203658 |
| Go_biological_process_2021 | Neuron projection extension (GO:1990138) | 0.004942017 | 0.012579679 | 2.306095783 |
| Go_biological_process_2021 | Neuron migration (GO:0001764) | 0.007481552 | 0.015517294 | 2.12600828 |
| Go_biological_process_2021 | Cell morphogenesis involved in neuron differentiation (GO:0048667) | 0.01135719 | 0.018272071 | 1.944729102 |
| Go_biological_process_2021 | Axon development (GO:0061564) | 0.011803729 | 0.018272071 | 1.927980765 |
| Go_biological_process_2021 | Neuron projection morphogenesis (GO:0048812) | 0.020854225 | 0.025927878 | 1.680805954 |
| Go_cellular_component_2021 | Apical dendrite (GO:0097440) | 0.001499295 | 0.025488011 | 2.824112974 |
| Go_cellular_component_2021 | Microtubule (GO:0005874) | 0.027053488 | 0.066704834 | 1.567776727 |
| Go_cellular_component_2021 | Axon (GO:0030424) | 0.030290255 | 0.066704834 | 1.518697074 |
| Go_cellular_component_2021 | Dendrite (GO:0030425) | 0.03995747 | 0.066704834 | 1.398402018 |
| Go_cellular_component_2021 | Microtubule cytoskeleton (GO:0015630) | 0.048834996 | 0.066704834 | 1.311268845 |

**Table S3. Neural related genes downregulated in neuron progenitor g, related to Fig. 3F.**

| Gene_set | P-value | Adjusted p-value | Logfoldchanges |
|---|---|---|---|
| *MINK1* | 0.001948794 | 0.027283114 | 2.710234094 |
| *MTOR* | 0.011654898 | 0.059656722 | 1.933491533 |
| *PRKG1* | 0.015816533 | 0.059656722 | 1.800888717 |
| *CSNK1E* | 0.019078183 | 0.059656722 | 1.719462988 |
| *PRKDC* | 0.02675888 | 0.059656722 | 1.572532063 |
| *MAPK3* | 0.0299963 | 0.059656722 | 1.522932307 |
| *MAPK8* | 0.033373192 | 0.059656722 | 1.476602255 |
| *PRKCB* | 0.037034828 | 0.059656722 | 1.431389672 |
| *SRC* | 0.03835075 | 0.059656722 | 1.416226143 |
| *MAPK1* | 0.048109397 | 0.062338273 | 1.317770082 |

**Table S4. Neuronal related GO enriched pathways downregulated in neuron progenitor g neuron cluster, related to Fig. 3G.**

| Gene_set | Term | P-value | Adjusted P-value | Genes | logfold changes |
|---|---|---|---|---|---|
| GO_Biological_Process_2021 | mitochondrial electron transport, NADH to ubiquinone (GO:0006120) | 0.005838817 | 0.01397254 | NDUFB2 | 2.233675133 |
| GO_Biological_Process_2021 | mitochondrial respiratory chain complex I assembly (GO:0032981) | 0.008675133 | 0.016193581 | NDUFB2 | 2.061723883 |
| GO_Biological_Process_2021 | mitochondrial ATP synthesis coupled electron transport (GO:0042775) | 0.0106126 6 | 0.018272071 | NDUFB2 | 1.974175757 |
| GO_Biological_Process_2021 | mitochondrial respiratory chain complex assembly (GO:0033108) | 0.013439889 | 0.018815844 | NDUFB2 | 1.871604329 |
| GO_Cellular_Component_2021 | mitochondrial respiratory chain complex I (GO:0005747) | 0.006287015 | 0.03562642 | NDUFB2 | 2.201555487 |
| GO_Cellular_Component_2021 | mitochondrial inner membrane (GO:0005743) | 0.048399681 | 0.066704834 | NDUFB2 | 1.3151575 |
| Reactome_2016 | Complex I biogenesis Homo sapiens R-HSA-6799198 | 0.007332288 | 0.029538174 | NDUFB2 | 2.13476051 |
| Reactome_2016 | Respiratory electron transport Homo sapiens R-HSA-611105 | 0.01314254 | 0.029538174 | NDUFB2 | 1.881320709 |
| Reactome_2016 | The citric acid (TCA) cycle and respiratory electron transport Homo sapiens R-HSA-1428517 | 0.022775849 | 0.034163774 | NDUFB2 | 1.64252542 |

**Table S5. Mitochondrial related genes downregulated in neuron progenitor g, related to Fig. 3H.**

| Gene_set | Logfoldchanges |
|---|---|
| *MAP1B* | 2.071300268 |
| *NDUFB2* | 1.701113224 |
| *TMSB4X* | 1.571030855 |

**Table S6. Neuronal related GO enriched pathways upregulated in neuron progenitor g neuron cluster after metformin treatment, related to Fig. 4E.**

| Gene_set | Term | Overlap | P-value | Fold change | Adjusted P-value |
|---|---|---|---|---|---|
| GO_Biological_Process_2021 | cell morphogenesis involved in neuron differentiation (GO:0048667) | 1/76 | 0,041025 | 1,386955 | 0,065871 |
| GO_Biological_Process_2021 | regulation of stem cell differentiation (GO:2000736) | 1/91 | 0,048938 | 1,310349 | 0,068338 |
| GO_Biological_Process_2021 | neuron projection morphogenesis (GO:0048812) | 1/140 | 0,074378 | 1,128553 | 0,087338 |
| GO_Biological_Process_2021 | axonogenesis (GO:0007409) | 1/240 | 0,124387 | 0,905224 | 0,133889 |
| GO_Cellular_Component_2021 | glutamatergic synapse (GO:0098978) | 1/69 | 0,037311 | 1,428163 | 0,087154 |
| GO_Cellular_Component_2021 | actin filament (GO:0005884) | 1/72 | 0,038904 | 1,410004 | 0,087154 |
| GO_Cellular_Component_2021 | neuron projection (GO:0043005) | 1/556 | 0,266706 | 0,573967 | 0,266706 |
| Reactome_2016 | RHO GTPases Activate WASPs and WAVEs Homo sapiens R-HSA-5663213 | 1/36 | 0,019627 | 1,707136 | 0,07048 |
| Reactome_2016 | Interleukin-1 signaling Homo sapiens R-HSA-446652 | 1/44 | 0,023941 | 1,620852 | 0,075287 |
| Reactome_2016 | Axon guidance Homo sapiens R-HSA-422475 | 2/515 | 0,031207 | 1,505745 | 0,075287 |
| Reactome_2016 | NCAM signaling for neurite out-growth Homo sapiens R-HSA-375165 | 1/266 | 0,136981 | 0,86334 | 0,160715 |
| Reactome_2016 | MAPK family signaling cascades Homo sapiens R-HSA-5683057 | 1/284 | 0,145603 | 0,836831 | 0,165434 |
| Reactome_2016 | Signalling by NGF Homo sapiens R-HSA-166520 | 1/450 | 0,221503 | 0,654621 | 0,223388 |
| Reactome_2016 | Signaling by GPCR Homo sapiens R-HSA-372790 | 1/1293 | 0,520671 | 0,283437 | 0,522877 |

**Table S7. Mitochondrial related GO enriched pathways upregulated in neuron progenitor g neuron cluster after metformin treatment, related to Fig. 4F.**

| Gene_set | Term | Overlap | P-value | Fold change | Adjusted P-value |
|---|---|---|---|---|---|
| GO_Biological_Process_2021 | DNA damage response, signal transduction by p53 class mediator resulting in transcription of p21 class mediator (GO:0006978) | 1/10 | 0,005488 | 2,26062 | 0,02933 |
| GO_Biological_Process_2021 | cellular iron ion homeostasis (GO:0006879) | 1/58 | 0,031449 | 1,502393 | 0,064139 |
| GO_Biological_Process_2021 | positive regulation of cellular amide +Metforminabolic process (GO:0034250) | 1/81 | 0,043669 | 1,359825 | 0,065871 |
| GO_Biological_Process_2021 | positive regulation of cellular protein +Metforminabolic process (GO:0032270) | 1/102 | 0,054704 | 1,261979 | 0,072544 |
| GO_Molecular_Function_2021 | ubiquitin protein ligase binding (GO:0031625) | 2/265 | 0,00889 | 2,051103 | 0,020036 |
| GO_Molecular_Function_2021 | protein kinase binding (GO:0019901) | 1/506 | 0,24569 | 0,609613 | 0,24569 |
| Reactome_2016 | +Metforminabolism of proteins Homo sapiens R-HSA-392499 | 3/1074 | 0,018418 | 1,734761 | 0,069286 |
| Reactome_2016 | Regulation of Apoptosis Homo sapiens R-HSA-169911 | 1/50 | 0,027165 | 1,565985 | 0,075287 |
| Reactome_2016 | Autodegradation of the E3 ubiquitin ligase COP1 Homo sapiens R-HSA-349425 | 1/51 | 0,027702 | 1,557493 | 0,075287 |
| Reactome_2016 | +Metforminabolism of amino acids and derivatives Homo sapiens R-HSA-71291 | 1/335 | 0,169608 | 0,770555 | 0,179942 |

**Table S8. Mitochondrial related GO enriched pathways upregulated in neuron progenitor g neuron cluster after metformin treatment, related to Fig. 4F.**

| Gene-set | scores | logfoldchanges | pvals | pvals_adj |
|---|---|---|---|---|
| *TUBA1A* | 9,039074 | 0,826918 | 1,58E-19 | 2,43E-15 |
| *ACTB* | 5,217101 | 0,438567 | 1,82E-07 | 0,000165 |
| *TUBA1B* | 4,97842 | 0,953523 | 6,41E-07 | 0,00047 |
| *H3F3B* | 4,188977 | 0,520429 | 2,8E-05 | 0,010036 |
| *PSMA7* | 6,673311 | 0,685721 | 2,5E-11 | 1,1E-07 |
| *MT-ATP6* | 4,256732 | 0,379086 | 2,07E-05 | 0,008087 |
| *MT-ND4* | 3,849178 | 0,213943 | 0,000119 | 0,033489 |
